# Compartmentalized localization of perinuclear proteins within germ granules in *C. elegans*

**DOI:** 10.1101/2024.03.25.586584

**Authors:** Xiaona Huang, Xuezhu Feng, Yong-hong Yan, Demin Xu, Ke Wang, Chengming Zhu, Meng-qiu Dong, Xinya Huang, Shouhong Guang, Xiangyang Chen

## Abstract

Biomolecular condensates are nonmembrane-enclosed organelles that play important roles in distinct biological processes. Germ granules are RNA-rich condensates that are often docked on the cytoplasmic surface of germline nuclei and exist in a variety of organisms. The *C. elegans* germ granule is a condensate that is subcompartmentalized into at least 6 distinct regions, including the P granule, Z granule, E granule, Mutator foci, SIMR foci and P-body. Although a number of perinuclear proteins have been found to be enriched in germ granules, their precise localization within the subcompartments of germ granules is still unclear. Here, we systematically labeled perinuclear localized proteins with fluorescent proteins via CRISPR/Cas9 technology and further explored the perinuclear localization of these proteins. Using this nematode strain library, we identified a series of proteins localized in Z or E granules. We found that proteins that participate in distinct piRNA processing steps were localized in particular subcompartments, including the P-, Z-, and E-granules. We further identified a novel subcompartment, the D granule, which was enriched between the P granule and the nuclear pore complex. Finally, we analyzed the perinuclear localization of these germ granule subcompartments in *mip-1/eggd-1* mutants and found that the LOTUS domain protein MIP-1/EDDG-1 was required for the establishment of the multiphase architecture of germ granules. Overall, our work revealed germ granule architecture and redefined the localization of perinuclear proteins within germ granules. Additionally, the library of genetically modified nematode strains expressing peri-nuclear proteins labeled with fluorescent tags will facilitate research on germ granules in *C. elegans*.

## Introduction

Biomolecular condensates are nonmembrane-enclosed organelles that consist of RNAs and proteins, the formation of which is likely elicited by phase separation and mediated by weak and multivalent interactions between RNA, intrinsically disordered proteins, and RNA-binding proteins^1–3^. Common biomolecular condensates include nucleoli, processing (P) bodies, Cajal bodies, stress granules and germ granules. The molecular constituents within these liquid droplet-like condensates exchange rapidly with the surrounding cellular contents^4,5^. The current model posits that these biomolecular condensates spatiotemporally bring proteins and RNA molecules together to orchestrate complicated RNA processing steps and coordinate gene expression^4,6^.

Many biomolecular condensates contain distinct immiscible condensed subcompartments, giving rise to multilayered liquid droplets that may facilitate sequential RNA processing reactions in a variety of RNP bodies^7–9^. For example, the nucleoli usually contain subcompartmentalized liquid phases composed of at least three distinct and coexisting subcompartments termed the fibrillar center (FC), dense fibrillar component (DFC), and granular component (GC), which are spatially organized, forming layered droplet organization^8,10^. The layered, multiphase droplet nature of nucleoli is thought to facilitate assembly line processing of ribosomal RNA (rRNA)^8^. Nascent rRNA transcripts undergo sequential processing steps by enzymes that localize to distinct intranucleolar subcompartments, before being exported from the nucleolus to the cytoplasm for final ribosome assembly. Recently, high-resolution fluorescence microscopy assays of fluorescent tag-labeled proteins enabled the investigation of the multilayered organization of biomolecular condensates directly in their cellular context and revealed the multiphasic condensate composition and spatial organization of many condensates, such as nucleoli, nuclear speckles, paraspeckles, stress granules and germ granules^11–16^. Although the underlying molecular mechanisms are largely unknown, the spatial organization of biomolecules into distinct subcompartments within biomolecular condensates may add another layer of internal composition regulation and play a fundamental role in facilitating their complex biological functions.

Germ granules are RNA-rich biomolecular condensates that are often docked on the cytoplasmic surface of animal germ cell nuclei and likely promote germ cell development and function by acting as organizational hubs for post-transcriptional gene regulation^17–21^. Germ granules are widely present in the germ cells of a variety of animals including worms, flies, zebrafish, Xenopus and mice, and are spatially organized into multilayered structures^21–25^. In *C. elegans*, recent studies have suggested that the germ granule is subcompartmentalized into several distinct regions that enclose particular sets of proteins, including P granule, Z granule, E granule, Mutator foci, SIMR foci and P-bodies^21,26,27^. The P granule was the first identified membraneless organelle to form via LLPS, which exhibits multiple liquid-like behaviors and has emerged as a leading model for the study of biomolecular condensates^28,29^. PGL-1 is a germline-expressed protein that is a constitutive protein component and is widely used as a marker protein of P granules^30^. The Z granule contains ZNFX-1 and WAGO-4, which promote the transgenerational inheritance of RNAi^14,31,32^. The E granule and Mutator foci are two independent subcompartments required for 22G RNA generation using a largely nonoverlapping set of RNA transcripts as templates^27,33,34^. The SIMR-1 foci, which are marked by SIMR-1, promotes siRNA amplification from piRNA targets and drive small RNA specificity for the nuclear Argonaute protein HRDE-1^35,36^. P-bodies, marked by CGH-1, are cellular aggregates of translationally repressed mRNPs that usually degrade mRNAs and inhibit their translation^26,37,38^. Remarkably, these immiscible germ granule subcompartments are not randomly ordered with respect to each other. For instance, many germ granules contain a single Z granule sandwiched between a P granule and a Mutator foci, forming ordered tri-condensate assemblages termed PZM granules^14^. Intriguingly, a recent study discovered a toroidal P granule morphology in the mid- and late pachytene regions of the germline, which encircles the other germ granule compartments in a consistent exterior-to-interior spatial organization, further revealing the hierarchical organization of germ granules^39^. However, little is known about how and why germ granules are divided into so many granular subcompartments and whether additional subcompartments of germ granules await discovery.

The multiphasic *C. elegans* germ granule provides multiple unique subcompartments to organize these perinuclear proteins and establish highly sophisticated perinuclear gene regulation networks, including small RNA-based gene regulatory pathways, which help to maintain germ cell totipotency and promote fertility^21,23,40^. For instance, mutator factors, including MUT-2/RDE-3, MUT-7, MUT- 8/RDE-2, MUT-14, MUT-15/RDE-5, MUT-16/RDE-6 and RDE-8, accumulate at the Mutator foci, contributing to siRNA amplification from poly(UG)-tailed RNA templates^34,41,42^. ZNFX-1, WAGO-4, PID-2/ZSP-1 and LOTR-1 accumulate in the Z granule, promoting RNAi inheritance and piRNA-induced gene silencing^14,15,43,44^. Approximately 90 *C. elegans* proteins have been found to be enriched in germ granules through a variety of methods, including proteomic approaches, such as TurboID/MS and IP/MS technologies^21,27,36,45,46^. Although high-resolution imaging and genetic analysis help us determine the sub-organelle localization of some perinuclear proteins, the precise localization of a considerable portion of these proteins within germ granules is still unclear. For instance, DDX-19, a conserved mRNA export factor whose position within germ granule is unknown, was reported to concentrate between the zones of PGL-1 and NPP-9, forming a tripartite architecture^47^. Deciphering the sub-organelle positioning of perinuclear proteins may help to understand perinuclear RNA processing and gene regulation networks.

Here, we systematically labeled reported perinucleus-localized proteins with fluorescent tags via CRISPR/Cas9 technology. We re-explored the subcellular localization of these proteins within germ granules and corrected some ambiguous and missing annotations in previous research. We found that proteins participating in distinct piRNA processing steps were each localized in particular subcompartments. We identified a novel subcompartment, named the D granule, that localized between the P granule and the nuclear pore complex. Furthermore, analysis of the formation of these germ granule compartments suggested that the architecture of the germ granule was maintained by the LOTUS domain protein MIP-1/EGGD-1. Overall, our work revealed the architecture of the multilayered germ granule and provided a resource for investigating comprehensive expression maps and functions of perinuclear proteins in *C. elegans*.

## Results

### Systematic labeling perinuclear proteins with fluorescence tags via CRISPR/Cas9 technology

The current model posits that *C. elegans* germ granule are divided into at least 6 subcompartments, including P granule, Z granule, E granule, Mutator foci, SIMR foci and P-bodies ^21,26,27^ (Fig. 1A). These subcompartments, in which distinct sets of proteins have been identified, are assembled in an orderly manner outside the nuclear envelope^21^. Additionally, in this study, we identified a novel subcompartment, the D granule (see Fig. 3 for details) (Fig. 1A). Particular proteins were used as marker proteins to visualize each subcompartment (Fig. 1B).

**Figure 1.**
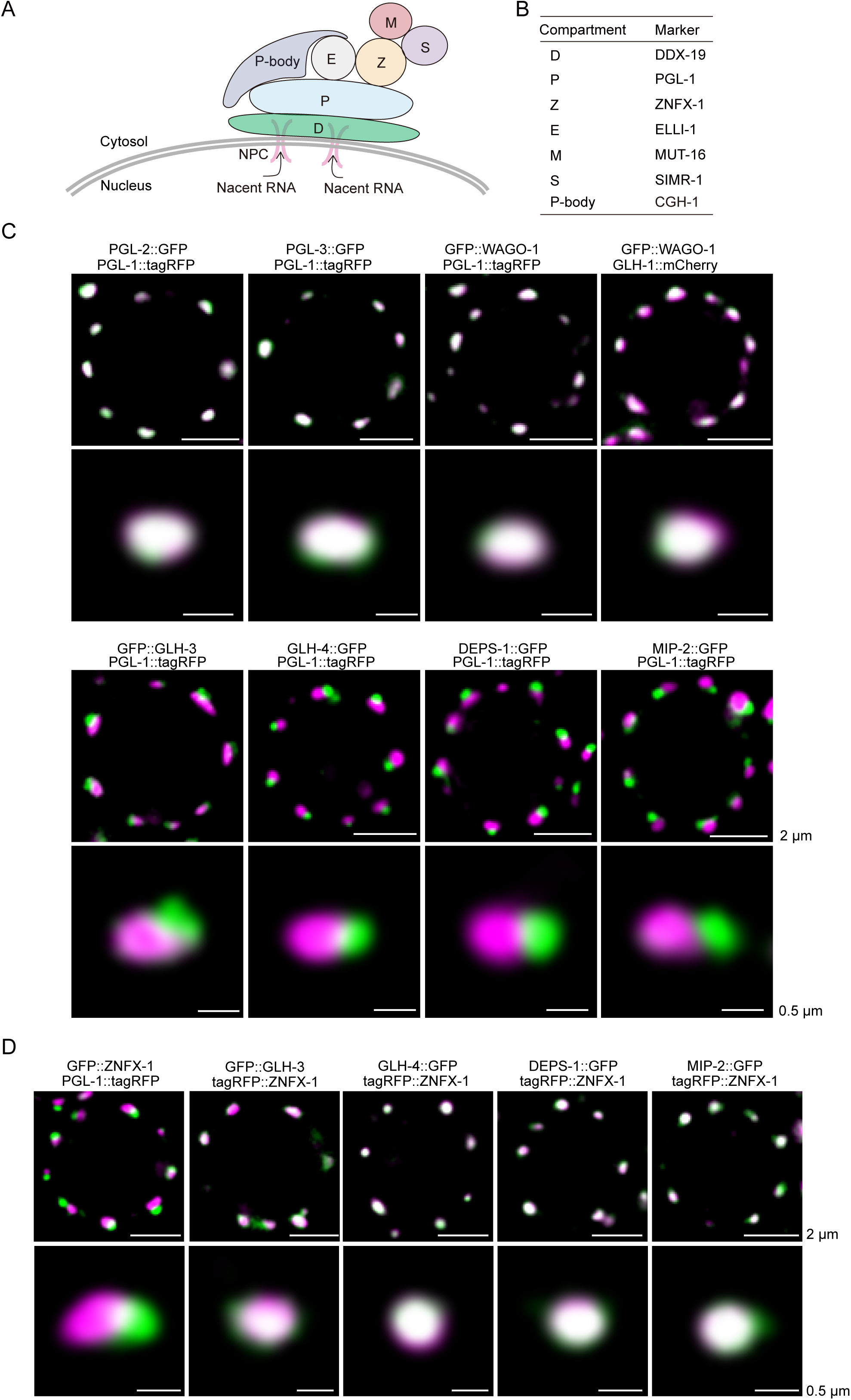
Identification of Z granule localized perinuclear proteins. (A) The model of germ granule architecture in the *C. elegans* germline. (B) Summary of the markers of each germ granule subcompartment. (C, D) Fluorescence micrographs of pachytene germ cells that express indicated fluorescent proteins. GLH-3, GLH-4, DEPS-1 and MIP-2 were enriched in the Z granule. Images were acquired with a Leica THUNDER Imaging System and deconvoluted using Leica Application Suite X software (version 3.7.4.23463). Images are representative of more than three animals.

More than half of the perinuclear proteins were previously annotated to be enriched in the P granule^21^. However, improvements in microscopy resolution and the identification of novel subcompartments prompted us to reinvestigate the subcellular localization of the reported perinuclear proteins within germ granules. Thus, we systematically tagged these proteins with fluorescent tags via CRISPR/Cas9-mediated knock-in technology (Fig. S1A) (see the methods section for details).

In this study, we have successfully tagged approximately 40 proteins with GFP, mCherry or tagRFP tags. The insertion of fluorescent tags was confirmed via DNA sequencing. As most of these genes are required for fertility^21^, we assessed the brood size of the strains and did not find significant alterations in most of the strains, compared to that of wild-type N2 worms (Fig. S1B), suggesting that the fluorescence-tagged proteins largely recapitulated their endogenous functions. We further collected published nematode strains expressing fluorescence-tagged perinuclear proteins, which were constructed previously in our laboratory^27,48^. In total, we collected a nematode strain library that consisted of more than 70 strains covering approximately 60 genes (Table S1).

We then crossed these tagged strains with each marker strain (Fig. 1B) and examined their perinuclear localization within germ granules. It should be noted that most of these proteins localize to both the cytosol and germ granule in germ cells or embryos, especially the P-body proteins, yet we mainly focused on deciphering the localization of these proteins within perinuclear germ granules in adult germ cells in this study. The localization of most of these proteins was consistent with the literature^21,49^ (Fig. S2A-S2D), although several proteins, such as MINA-1, PAB-1, PLP-1 and GLD-4, showed poor accumulation within germ granules, making it difficult to clarify their intragranular positioning (Fig. S2E). However, we did identify a considerable number of proteins that accumulated in other compartments of the germ granule rather than in the P granule (see below for details). The results are summarized in Table 1.

**Table 1:**
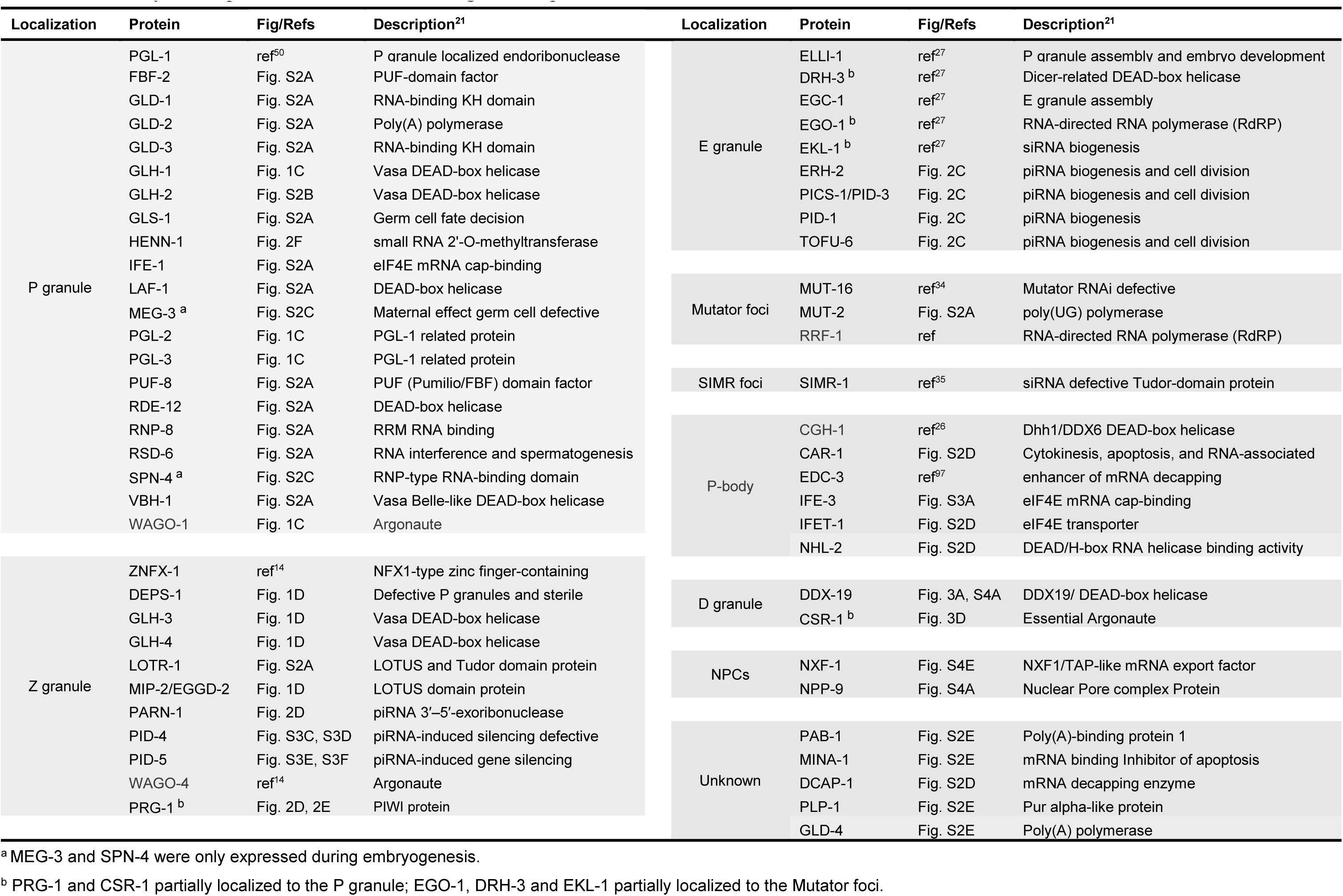
Summary of the compartmentalized localization of germline proteins within germ granules.

### GLH-3, GLH-4, DEPS-1 and MIP-2 were enriched in the Z granule

We first examined the localization of common P granule markers and the factors required for P granule assembly. PGL-1::tagRFP was used as the marker of P granules^50^. The PGL family proteins, including PGL-1, PGL-2 and PGL-3, localize in germ granules and function redundantly in *C. elegans* germline development^51^. As reported, PGL-2 and PGL-3 colocalized with PGL-1 in the P granule (Fig. 1C). WAGO-1, an Argonaute protein expressed in the germline and involved in the RNAi pathway, also colocalized with PGL-1::tagRFP, as previously reported (Fig. 1C)^52^.

Vasa DEAD-box helicases are widespread markers of germ cells among different species^53^. *C. elegans* has four Vasa family members, GLH-1, GLH-2, GLH-3 and GLH-4, all of which are associated with germ granules^54^. Among these proteins, GLH-1 and GLH-4 are required for proper association of the PGL family proteins with P granules, and for the formation and organization of germ granules^54–56^. We examined whether these proteins are localized in the P granule. GLH-1 colocalized with WAGO-1 (Fig. 1C), and GLH-2 colocalized with PGL-1 (Fig. S2B), suggesting that GLH-1 and GLH-2 accumulated in the P granule. However, GLH-3 and GLH-4 did not colocalize with PGL-1, but colocalized with tagRFP::ZNFX-1, suggesting that GLH-3 and GLH-4 preferentially accumulated in the Z granule (Figs. 1C and 1D).

DEPS-1 is required for the assembly of P granules, the loss of which disrupts P-granule structure and function^57^. DEPS-1 is required for the accumulation of *glh-1* and *rde-4* mRNAs to promote the RNAi response, germ cell proliferation and fertility at elevated temperature^57,58^. DEPS-1 did not colocalize with PGL-1::tagRFP (Fig. 1C). However, similar to GLH-4, DEPS-1 colocalized with ZNFX-1 and accumulated in Z granules (Fig. 1D).

MIP-1/EGGD-1 and MIP-2/EGGD-2 are two LOTUS domain-containing proteins that recruit *C. elegans* Vasa to germ granules^45,46^. Animals lacking MIP-1/EGGD-1 and MIP-2/EGGD-2 exhibit P granule detachment from the nuclear envelope, temperature-sensitive embryonic lethality and sterility^45,46^. MIP-1/EGGD-1 is reported to be enriched in P granules, yet the cellular location of MIP-2 in germ cells is still unclear^45,46^. We found that MIP-2/EGGD-2 was enriched in Z granules, rather than in P granules in germ cells (Figs. 1C and 1D). LOTR-1, another LOTUS domain-containing protein, colocalized with tagRFP::ZNFX-1 and accumulated in the Z granule, as reported previously (Fig. S2A)^44^.

Overall, we found that a number of factors, including GLH-3, GLH-4, DEPS-1 and MIP-2, which were previously annotated to localize in the P granule, were likely components of the Z granule.

### piRNA processing and maturation factors accumulate in distinct germline subcompartments

In *C. elegans*, many of the factors involved in the biogenesis of small regulatory RNAs, including siRNAs and piRNAs, as well as the Argonaute (AGO) proteins that bind these small RNAs, are largely enriched in germ granules^21^. We and other groups previously showed that the factors localized in E granules and Mutator foci contribute to the production of distinct cohorts of siRNAs using non-overlapping RNA templates^27,33,34^. piRNAs, which are 21 nt in length and start with 5′-monophosphorylated uracil, are produced from Pol II-transcribed piRNA precursors, exported to the cytoplasm, and subjected to a series of processing steps in the germline (Fig. 2A)^59–61^.

**Figure 2.**
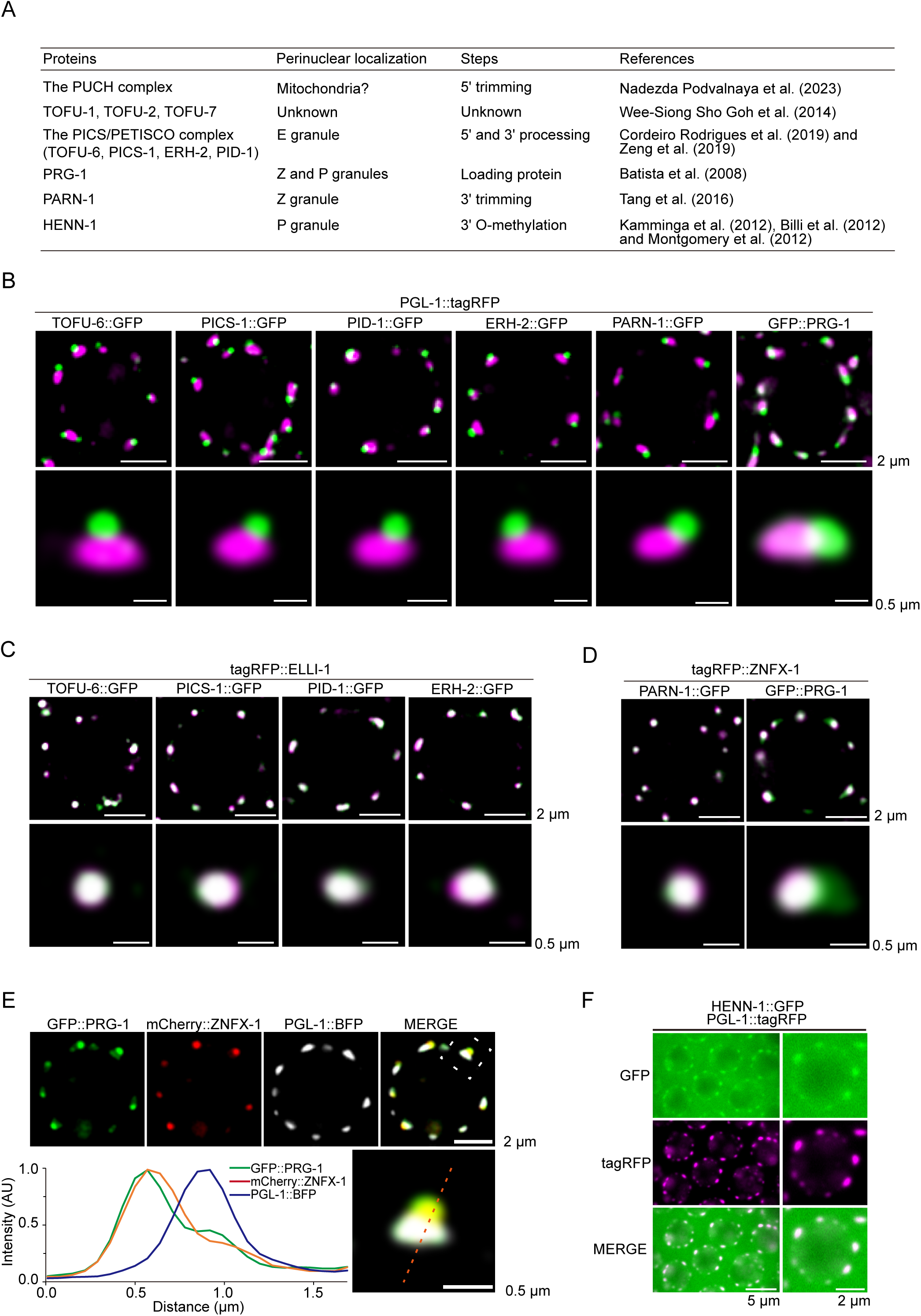
Subcellular localization of piRNA processing factors in *C. elegans*. (A) Summary of proteins participating in the processing of piRNA precursors in *C. elegans*. (B) Fluorescence micrographs of pachytene germ cells that express PGL-1::tagRFP and the indicated GFP-tagged proteins. The PICS/PETISCO complex did not colocalize with PGL-1. PRG-1 partially colocalized with PGL-1. (C) Fluorescence micrographs of pachytene germ cells expressing tagRFP::ELLI-1 and the indicated GFP-tagged proteins. The PICS/PETISCO complex was enriched in the E granule. (D) Fluorescence micrographs of pachytene germ cells expressing tagRFP::ZNFX-1 and the indicated GFP-tagged proteins. PARN-1 accumulated in the Z granule, and PRG-1 was largely enriched in the Z granule. (E) Fluorescence micrographs of pachytene germ cells from animals expressing GFP::PRG-1, mCherry::ZNFX-1 and PGL-1::BFP. The intensity of fluorescence along the dotted line was calculated by Leica Application Suite X software (version 3.7.4.23463). (F) Pachytene germ cells of animals that express HENN-1::GFP and PGL-1::tagRFP. All images are representative of more than three animals.

The PICS/PETISCO complex consists of four core subunits, namely, TOFU-6, PICS-1/PID-3, ERH-2 and PID-1^48,62^. The PICS/PETISCO complex stabilizes the PUCH complex and facilitates 5*’* trimming of piRNA precursors^61^. We previously reported that the knocking down *csr-1* and *glh-1*, two reported P granule components, failed to disrupt the perinuclear PICS foci, implying that the PICS/PETISCO complex may not localize to the P granule^48^. Consistent with this idea, these proteins did not colocalize but were rather adjacent to the P granule marker PGL-1 (Fig. 2B). We observed that the GFP-tagged PICS complex components, including TOFU-6, PICS-1/PID-3, ERH-2 and PID-1, were all colocalized with tagRFP::ELLI-1, which is a marker of the E granule (Fig. 2C). Intriguingly, IFE-3, a *C. elegans* eIF4E homolog that interacts with PID-1 and likely involves in 5*’* maturation of piRNA precursors, seems to accumulate in the P-body (Fig. S3A).

*parn-1* encodes a conserved 3*’* to 5*’* RNA exoribonuclease, the loss of which elicits the accumulation of untrimmed piRNA precursors with 3*’* extension^63^. PARN-1 did not colocalize with PGL-1, but rather colocalized with ZNFX-1, suggesting that PARN-1 accumulates in the Z granule (Figs. 2B and 2D). PID-2/ZSP-1 is exclusively required for the proper assembly of Z granules^15,43^. In *pid-2* mutants, PARN-1 remained colocalized with ZNFX-1, similar to that observed for GLH-3, GLH-4 and MIP-2 (Fig. S3B).

PRG-1 is a member of the *C. elegans* PIWI family that exclusively binds to piRNAs and mediates the genome-wide surveillance of germline transcripts^64–67^. We found that PRG-1 accumulated in both the P and Z granules and was largely enriched in the Z granule (Figs. 2B and 2D). Tricolor images from animals simultaneously expressing PGL-1::BFP, mCherry::ZNFX-1 and PRG-1::GFP further support the above conclusion (Fig. 2E). Additionally, PRG-1 remained colocalized with ZNFX-1 in *pid-2* mutants (Fig. S3B). HENN-1 is a 3’-2’-O-methyltransferase that methylates small RNA molecules including piRNAs and ERGO-1-bound 26G RNAs^68–70^. Consistent with previous reports, HENN-1 accumulated in the P granule (Fig. 2F)^69^.

PID-4 and PID-5 were identified as PID-2 interactors that act redundantly for piRNA sensor silencing, affect the size and appearance of Z granules, and regulate 22G RNA production^43^. Both PID-4 and PID-5 are required for germline immortality^43^. We found that both PID-4 and PID-5 did not colocalize with PGL-1, but largely colocalized with tagRFP::ZNFX-1 and largely accumulated in Z granules in germ cells of both L4 and adult animals, as previously reported (Fig. S3C-S3F)^43^.

As the piRNA processing and maturation factors were each enriched in distinct subcompartments of the germ granule, we speculated that it is very likely that the piRNA processing intermediates could be transported between different subcompartments and/or within the cytosol.

### DDX-19 and CSR-1 define a new subcompartment of the germ granule

DDX-19 is a conserved mRNA export factor that is reported to concentrate between the zones of PGL-1 and NPP-9 and therefore forms a tripartite architecture^47^. As reported, GFP::DDX-19 was enriched under the P granule and above the nuclear pore complex (as shown by NPP-9::mCherry) in germline cells (Figs. 3A, S4A and S4B). Therefore, we named the DDX-19 foci as D granules. Interestingly, D granules disintegrated during embryogenesis and DDX-19 directly colocalized with NPP-9 in the embryos, suggesting that the assemblage of D granules was developmentally regulated (Fig. S4C).

**Figure 3.**
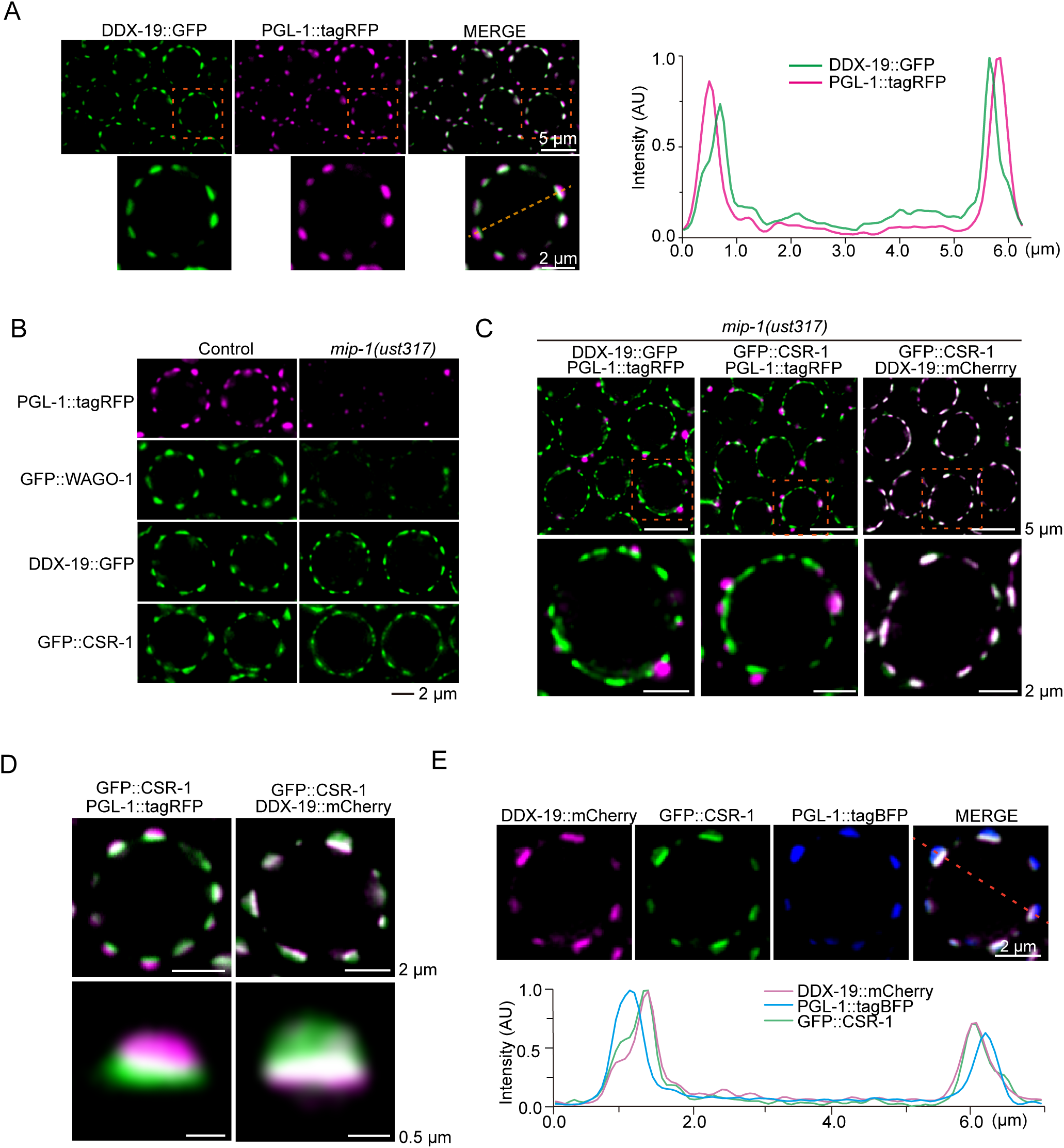
DDX-19 and CSR-1 are enriched in a novel germ granule subcompartment. (A) Left. Fluorescence micrographs of pachytene germ cells expressing DDX-19::GFP and PGL-1::tagRFP. Right, the fluorescence intensity along the dotted line, which was calculated by Leica Application Suite X software (version 3.7.4.23463). (B) Images of representative meiotic germ cells from the indicated animals. Loss of MIP-1/EGGD-1 does not block the perinuclear localization of DDX-19 or CSR-1. (C) Fluorescence micrographs of *mip-1/eggd-1* animals expressing the indicated fluorescent proteins. (D) Fluorescence micrographs of animals expressing GFP::CSR-1 and PGL-1::tagRFP, or GFP::CSR-1 and DDX-19::mCherry. (E) Up, pachytene germ cells of animals that express the indicated fluorescent proteins. Down, the fluorescense intensity along the dotted line, which was calculated by Leica Application Suite X software (version 3.7.4.23463). CSR-1 was mainly enriched in the D granule and moderately localized in the P granule. All images are representative of more than three animals.

The LOTUS domain protein MIP-1/EGGD-1 is required for the perinuclear localization of P granules, Z granules and Mutator foci in germ cells^71^. The loss of MIP-1/EGGD-1 disrupted the PGL-1- and WAGO-1-marked P granule (Fig. 3B), but did not affect the perinuclear localized DDX-19 foci, suggesting that MIP-1/EGGD-1 was not required for the perinuclear anchoring of the D granule (Fig. 3B). The residue PGL-1::tagRFP still localized adjacent to DDX-19::GFP in the absence of MIP-1/EGGD-1 (Fig. 3C).

CSR-1 is the unique essential Argonaute protein in *C. elegans*. The genomic *csr-1* locus encodes two isoforms, CSR-1a and CSR-1b, which vary only in their N-terminus^72,73^. CSR-1a is expressed during spermatogenesis and in several somatic tissues, including the intestine; CSR-1b is expressed constitutively in the germline^72,73^. CSR-1 largely accumulates to perinuclear granules upon the depletion of MIP-1/EGGD-1, as well as in *glh-1;glh-4* mutants ^56,71^. Here, we synchronously tagged two isoforms^74^, and focused on the cellular localization of CSR-1b in germ cells. We wondered if CSR-1, like DDX-19, accumulated in the D granule at the base of the P granule as well. Indeed, CSR-1 mainly colocalized with DDX-19 yet revealed a moderate localization in the P granule (Fig. 3D). Tricolor images from animals simultaneously expressing PGL-1::tagBFP, DDX-19::mCherry and GFP::CSR-1 further supported the above conclusion (Fig. 3E). The loss of MIP-1/EGGD-1 dramatically decreased the localization of CSR-1 in P granules but had a marginal, if any, effect on D granule localization (Figs. 3B, 3C, and S4D).

*C. elegans* NXF-1, the ortholog of the essential mRNA export factor NXF1/TAP in humans and Mex67p in yeast, is reported to concentrate below P granules, whose localization at the nuclear rim is very similar to that of NPP-9^47,75^. We found that NXF-1 mainly localized in the nucleus, as recently reported (Fig. S4E) ^76^. Additionally, a small portion of this protein was enriched at the base of DDX-19 condensates, suggesting that perinuclear NFX-1 may be enriched in the NPCs (Fig. S4E).

Overall, we identified a novel perinuclear D granule containing DDX-19, and CSR-1 and a tripartite architecture of the P granule, the D granule and the nuclear pore complex.

### The LOTUS domain protein MIP-1/EGGD-1 sculptures the architecture of perinuclear germ granules

Recent reports showed that the loss of MIP-1/EGGD-1 disrupted perinuclear positioning of P granules, Z granules and Mutator foci, and induced the mislocalization of these subcompartments within the adult gonad^45,46,71^. Here, we examined whether the LOTUS domain MIP-1/EGGD-1 is required for the assembly and perinuclear localization of other germ granule subcompartments. Consistently, the depletion of MIP-1/EGGD-1 resulted in the dispersal of perinuclear PGL-1, ZNFX-1 and MUT-16 and the accumulation of aggregates at the rachis (Figs. 4A and S5). Additionally, when MIP-1/EGGD-1 was deleted, the SIMR foci also largely detached from the perinuclear zone, and accumulated at the rachis (Fig. S5). However, the D granule (marked by DDX-19 and CSR-1), the E granule (marked by ELLI-1 and EGC-1) and the P-body (marked by CGH-1) still exhibited pronounced foci formation at the periphery of germ cell nuclei, suggesting that MIP-1/EGGD-1 was not required for the perinuclear localization of these three subcompartments (Figs. 4A and S5). Additionally, the loss of MIP-1/EGGD-1 did not induce the generation of dissociative D or E granules in the rachis. Conversely, the rachis-localized E granules (marked by ELLI-1 or EGC-1), which exist in wild-type animals, seemed to disappear in *mip-1/eggd-1* mutants (Figs. 4A and S5). Since P-body proteins were also enriched as irregularly shaped aggregates in the rachis of the germline in wild-type animals, it is unclear whether P-bodies were altered in the rachis upon the depletion of MIP-1/EGGD-1 (Fig. S5).

**Figure 4.**
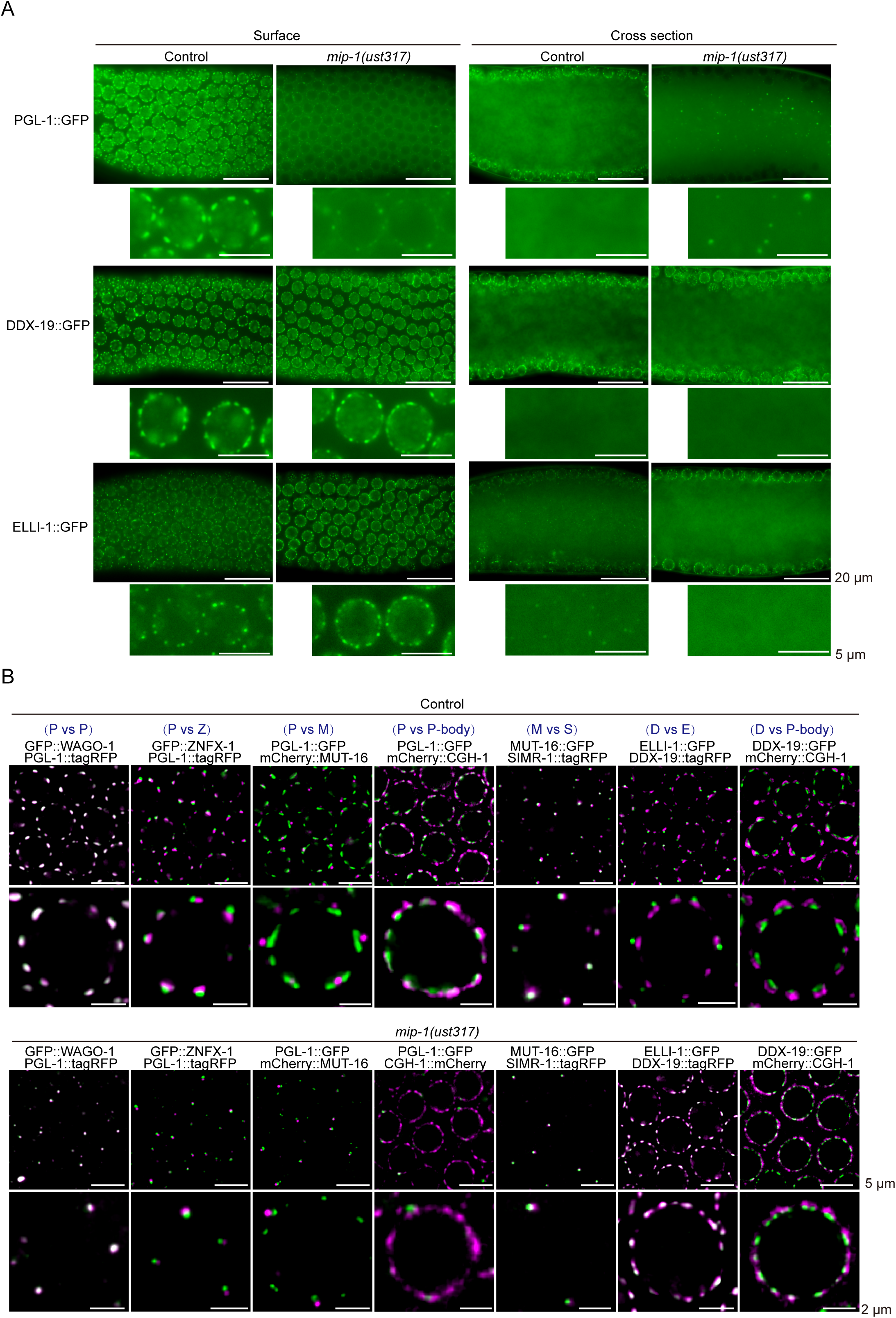
The LOTUS domain protein MIP-1/EGGD-1 is required for the organization of the germ granule architecture. (A) Fluorescence micrographs of the surface and rachis of the germline in live adult animals expressing the indicated proteins in wild-type and *mip-1/eggd-1* animals. Loss of MIP-1/EGGD-1 does not elicit the accumulation of DDX-19 or ELLI-1 foci in the rachis. (B) Fluorescence micrographs of wild-type and *mip-1/eggd-1* animals expressing indicated proteins. The D and E granules fused with each other upon the depletion of MIP-1/EGGD-1. All images are representative of more than three animals.

We further tested the colocalization of different germ granule subcompartments in *mip-1/eggd-1* mutants to examine whether these subcompartments remain immiscible. The residual PGL-1::tagRFP, GFP::ZNFX-1, mCherry::MUT-16 and SIMR-1::tagRFP remained separated from each other in *mip-1/eggd-1* mutants (Fig. 4B). Interestingly, the depletion of MIP-1/EGGD-1 induced the colocalization of ELLI-1::GFP and DDX-19::tagRFP, suggesting the disruption of immiscibility between the D and E granules and the likely fusion of these two condensates (Fig. 4B). The P-body marker CGH-1 still accumulated in the perinuclear region and did not fuse with D granules (Fig. 4B). Together, these data further suggested that the LOTUS domain protein MIP-1/EGGD-1 is involved in the organization of germ granule architecture.

## Discussion

Here, using CRISPR/Cas9 technology, we systemically tagged perinuclear proteins with fluorescent tags and examined their localization within germ granules. We imaged these perinuclear proteins and found that these proteins were enriched in specialized subcompartments of germ granules. Especially, proteins involved in particular piRNA processing steps were enclosed in different subcompartments. We also identified a novel subcompartment, termed the D granule, which consists of DDX-19 and CSR-1 and is located between the P granule and the nuclear pore complex. Finally, we investigated germ granule architecture in *mip-1/eggd-1* mutants and found that the LOTUS domain protein MIP-1/EGGD-1 was required for the immiscibility and multilayered organization of subcompartments within germ granules. Together, these data extend our understanding of the germ granule architecture, and the library of nematode strains expressing fluorescence-tagged perinuclear proteins may provide a powerful resource for further investigating the compartmentalization and function of germ granules.

Intracellular condensates are highly multicomponent systems that enclose a variety of RNP bodies in micron-scale membraneless organelles, undertaking diverse RNA processing events or sequential RNA processing reactions^7,13,77–79^. The compartmentalized structure of biomolecular condensates may provide an ingenious and efficient strategy for cells to control the spatial partitioning and processing of these molecules. For instance, the nucleolar proteome consists of hundreds of different proteins that are segregated into at least three distinct compartments of the nucleolus, including the FC, DFC and GC regions^8^. Cells control and coordinate successive ribosome production steps by generally allocating pre-rRNA transcription, rRNA processing and ribosome assembly events to the FC, DFC and GC compartments respectively^8^. Like the nucleolus, *C. elegans* germ granule consists of multiple subcompartments, including, but possibly not limited to the P, Z, E, M, S and D compartments and the P-body. However, it is still largely unknown why and how germ granules are compartmentalized into so many subcompartments. As a perinuclear organelle directly bound to the nuclear pore complex, it is likely that the germ granule is the principal site of mRNA export in germ cells to sort mRNAs to their destined regulatory molecules before being released into the cytoplasm^21,47,80^. The multiphase organization of the germ granule may provide a robust and efficient platform for enriching diverse regulatory molecules and establishing corresponding gene regulatory networks, especially the small RNA-based gene regulation pathways.

The capped piRNA precursors are transcribed from thousands of genomic loci on LG IV and go through sequential cytoplasmic processing steps before turning into mature piRNAs in *C. elegans*^59,81^. In this study, we found that factors involved in particular piRNA processing steps were enriched in distinct subcompartments within germ granules, including E granule-localized PICS/PETISCO complex, Z granule-localized PARN-1 and P granule-localized HENN-1. Additionally, the *C. elegans* piRNA-binding Argonaute protein PRG-1 localizes to both the Z and P granules and preferentially accumulates in the Z granule. The compartmental positioning of piRNA machineries within germ granules may help partition sequential piRNA processing steps in space and time, promoting piRNA maturation in a fluent and choreographed fashion. For instance, in mouse prospermatogonia (ProSg), pi-bodies and piP-bodies, which are often in close proximity to each other, facilitate the processing of primary and secondary pre-pachytene piRNAs, respectively, and promote the silencing of transposable elements^23,82^. Examination of piRNA biogenesis upon artificial modification of the subregional localization of piRNA processing factors within germ granules may help to decipher whether and how multiple immiscible condensed phases contribute to piRNA maturation.

RNAs transcribed in the nucleus should pass through nuclear pore complexes (NPCs) before being transported to the cytoplasm or the cocytoplasmic vesicles of the germline for translation^83^. Interestingly, it has been reported that approximately 75% of the nuclear pores in *C. elegans* germ cells are associated with germ granules, which therefore sequester large amounts of mRNAs^47,84^. It was suggested that the perinuclear germ granules are the main sites of mRNA output in adult *C. elegans* germ cells^47^. Since the D granule concentrates between the P granule and the NPCs, it might act as the first condensate to house the RNPs exported from the nucleus. Consistently, DDX-19 is a predicted DEAD-box helicase related to DDX19 in mammals or Dbp5p in yeast that interacts with the nucleoporin Nup159/CAN/Nup214 and functions in mRNA export from the nucleus through remodeling mRNPs on the cytoplasmic face of the nuclear pore^85–88^. Thus, the D granule may participate in sorting mRNA export from the nucleus to other subcompartments of germ granules. CSR-1 protects the germline transcriptome against the silencing activities of the piRNA genome surveillance pathway, further suggesting that the D compartments may be the first site for the fate determination of RNAs^89,90^. The biotin ligase, TurboID, which can robustly biotinylate proximal proteins of a target protein in living cells, has been widely applied for proteomic mapping of a wide range of protein complexes and cellular structures^46,91^. Using the TurboID–based proximity labeling system to decipher the D granule proteome may further help to define its biological functions. Moreover, developing new techniques to decipher the transcribed RNA molecules within these distinct subcompartments may elucidate whether and how germ granules are compartmentalized and regulated.

The mechanism by which condensates form and maintain internal subcompartments is unclear. Studies on the characteristic core-shell spheroidal structure of the nucleolus reveal that differences in subcompartment surface tension, which arises from the sequence features of macromolecular components, including RNAs and proteins, may determine the immiscibility of different subcompartments and the multilayered organization^10,92,93^. Similarly, these biological molecules are essential for the formation and hierarchical organization of germ granule subcompartments within *C. elegans* germ granules. For instance, the intrinsically disordered protein MUT-16 mediates the assembly of Mutator foci. The depletion of MUT-16 elicited the diffusion of Mutator foci components in the cytosol^34,41^. PID-2 is another intrinsically disordered protein that exclusively modulates Z granule morphology^15,43^. Additionally, the injection of young adult gonads with the transcriptional inhibitor a-amanitin or the depletion of NXF-1, an essential mRNA export factor in *C. elegans*, elicited the loss of PGL-1 foci and MUT-16 foci in pachytene cells, suggesting that RNA molecules are also required for the assembly of multiple condensates^47,94^. We and others found that the loss of MIP-1/EGGD-1 leads to extensive germ granule remodeling, including the separation of the P, Z, M and S compartments from the nuclear envelope and the fusion of the E and D compartments. How does the LOTUS domain protein MIP-1/EGGD-1 participate in the organization of germ granule subcompartments? The LOTUS domain, which is widely conserved in eukaryotes, has been identified within several proteins, such as Tudor domain-containing protein 5 (TDRD5), TDRD7, Oskar, and meiosis regulator and mRNA stability factor 1 (MARF1)^95,96^. All of these proteins play essential roles in animal development with a prominent function in gametogenesis. Interestingly, these proteins also emerged as key regulators of germ granule organization in multiple organisms. For example, TDRD-7 promotes the fusion of piP-bodies and early chromatoid bodies to form mature chromatoid bodies during spermatogenesis in mice^82^. In *Drosophila*, Oskar, which is the primary determinant of germline progenitors, has multiple roles in the organization of the germ plasm via the assembly and anchoring of germ plasm components at the egg cortex^20,22,24^. Intriguingly, LOTUS domains were reported to harbor both RNA binding activity, particularly for G-rich/G4 RNAs, and protein binding activity, especially for RNA helicases^95,96^. Here, we found that the loss of MIP-1/EGGD-1 destroyed *C. elegans* germ granules without affecting the perinuclear localization of D granules. Therefore, we speculate that MIP-1/EGGD-1 may participate in the allocation of RNA molecules from the D granule to other germ granule subcompartments, thereby resulting in the establishment of different surface tensions of distinct germ granule subcompartments to confer immiscibility and promote spatially organized architecture. Further studies are required to investigate the functions of the LOTUS domain proteins in the organization of germ granule architecture.

## Acknowledgments

We are grateful to Dr. Scott Kennedy and members of the Guang laboratory for their comments. We are grateful to Dr. Donglei Zhang for sharing unpublished results. We are grateful to the International *C. elegans* Gene Knockout Consortium and the National Bioresource Project for providing the strains. Some strains were provided by the CGC, which is funded by the NIH Office of Research Infrastructure Programs (P40 OD010440).

## Author contributions

X.H., X.C. and S.G. conceptualized the research; X.F., C.Z., X.H., X.C. and S.G. designed the research; X.C., X.H., X.F., Y.H., D.M., and W.K. performed the research; X.H. and X.F. contributed new reagents; X.C. and S.G. wrote the paper.

## Materials and Methods

### C. elegans strains

The Bristol strain N2 was used as the standard wild-type strain. All strains were grown at 20 °C unless otherwise specified. The strains used in this study are listed in Table S2.

### Construction of transgenic strains

The coding regions of *gfp::3xflag, 3xflag::gfp*, *mCherry* or *tagRFP* fused to a linker sequence (GGAGGTGGAGGTGGAGCT), were inserted upstream of the stop codon or downstream of the initiation start codon using the CRISPR/Cas9 system. Plasmids containing repair templates were generated using a ClonExpress MultiS One Step Cloning Kit (C113-02, Vazyme). The injection mixture contained pDD162 (50 ng/mL), a repair plasmid (50 ng/mL), a marker plasmid (pSG259(*myo-2p::gfp*::*unc-54utr*) or pSG280(*sqt-1p::sqt-1(e1350)::sqt-1utr*)) (5 ng/mL) and two or three gRNAs targeting sequences proximal to the N-termini or C-termini of the genes (each sgRNA plasmid, 20 ng/mL). The mixture was injected into adult animals. Two strategies were applied to identify targeted animals. (1) For knock-in lines using pSG259(*myo-2p::gfp*) as the co-injection marker, F1 worms expressing pharyngeal GFP were isolated under a Leica M165 FC fluorescence stereomicroscope and transferred onto individual NGM plates to lay F2 animals. The targeted animals with *gfp* insertions were screened by PCR. (2) For knock-in lines using pSG280(*sqt-1p::sqt-1(e1350)*) as the co-injection marker. Strategies were adjusted from an earlier study^1^. 8-10 F1 rollers were picked onto microscope slides, and the GFP fluorescence signals from germ cells were observed under a Leica DM4 B microscope. Worms with observable green fluorescence within the germline were transferred from the slides onto individual NGM plates to lay F2 worms. Then, 16 F2 adult worms were singled onto individual NGM plates, and the homozygous transgenes were subsequently identified by evaluating fluorescence signals in F3 animals followed by genotyping.

### Brood size

L3 worms were placed individually onto fresh NGM plates. The numbers of progeny that reached the L2 or L3 stage were scored. n=10 animals.

### Microscopy and imaging

To image larval stages, animals were immobilized in ddH_2_O with 0.5 M sodium azide and mounted on glass slides before imaging. To image germ cells in adult animals, worms were dissected in 2 µl of 0.4x M9 buffer with 0.1 M sodium azide on a coverslip and then mounted on freshly made 1.2%-1.4% agarose pads. The Leica TNUNDER Imaging System was used, equipped with a K5 sCMOS microscope camera and an HC PL APO 100x/1.40-0.70 oil objective. Images were taken and deconvoluted using Leica Application Suite X software (version 3.7.4.23463). Images in Figs. 2F, 4A, S2, S3C-3F, S4C and S5 were not deconvoluted. For the same proteins under different genetic backgrounds, equally normalized images were exported, and contrasts of images were equally adjusted between control and experimental sets. As the expression levels of different perinuclear proteins vary, the display values of fluorescence images showing the relative position of perinuclear proteins within germ granules were manually adjusted to visualize these proteins.

## Supplemental figure legends

**Figure S1.**
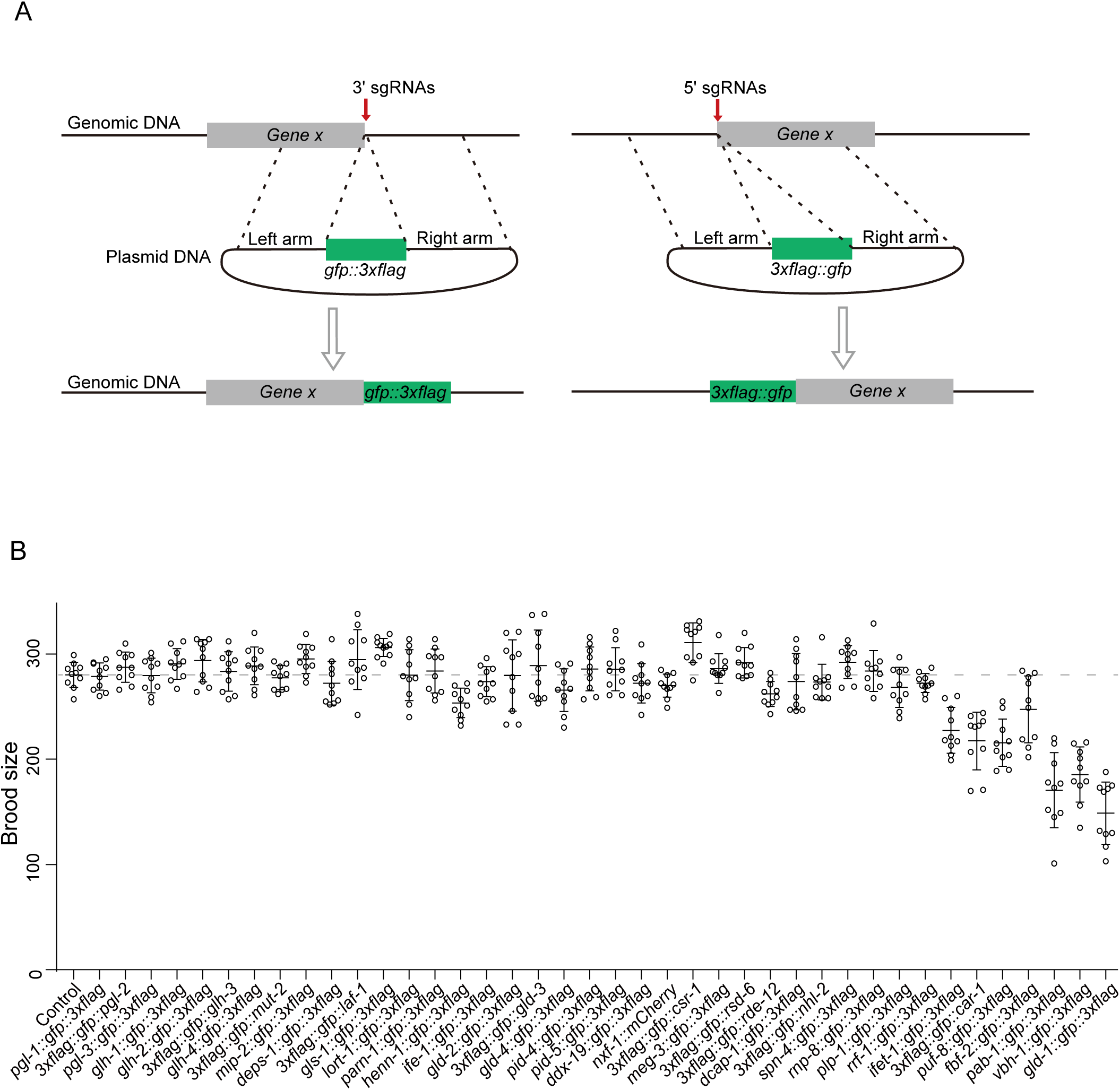
Schematic of gene editing events using CRISPR/Cas9 technology. (A) Schematic of the insertion of DNA elements encoding fluorescent proteins into the designed genomic loci using CRISPR/Cas9 technology. The coding regions of *gfp::3xflag* and *3xflag::gfp*, which were fused to a linker sequence, were inserted upstream of the stop codon or downstream of the initiation start codon respectively. (B) Brood sizes of the indicated animals at 20 °C. The bleached embryos were hatched and grown at 20 °C. Then, L3 worms were transferred individually onto fresh NGM plates. The number of progeny worms was scored. n=10 animals.

**Figure S2.**
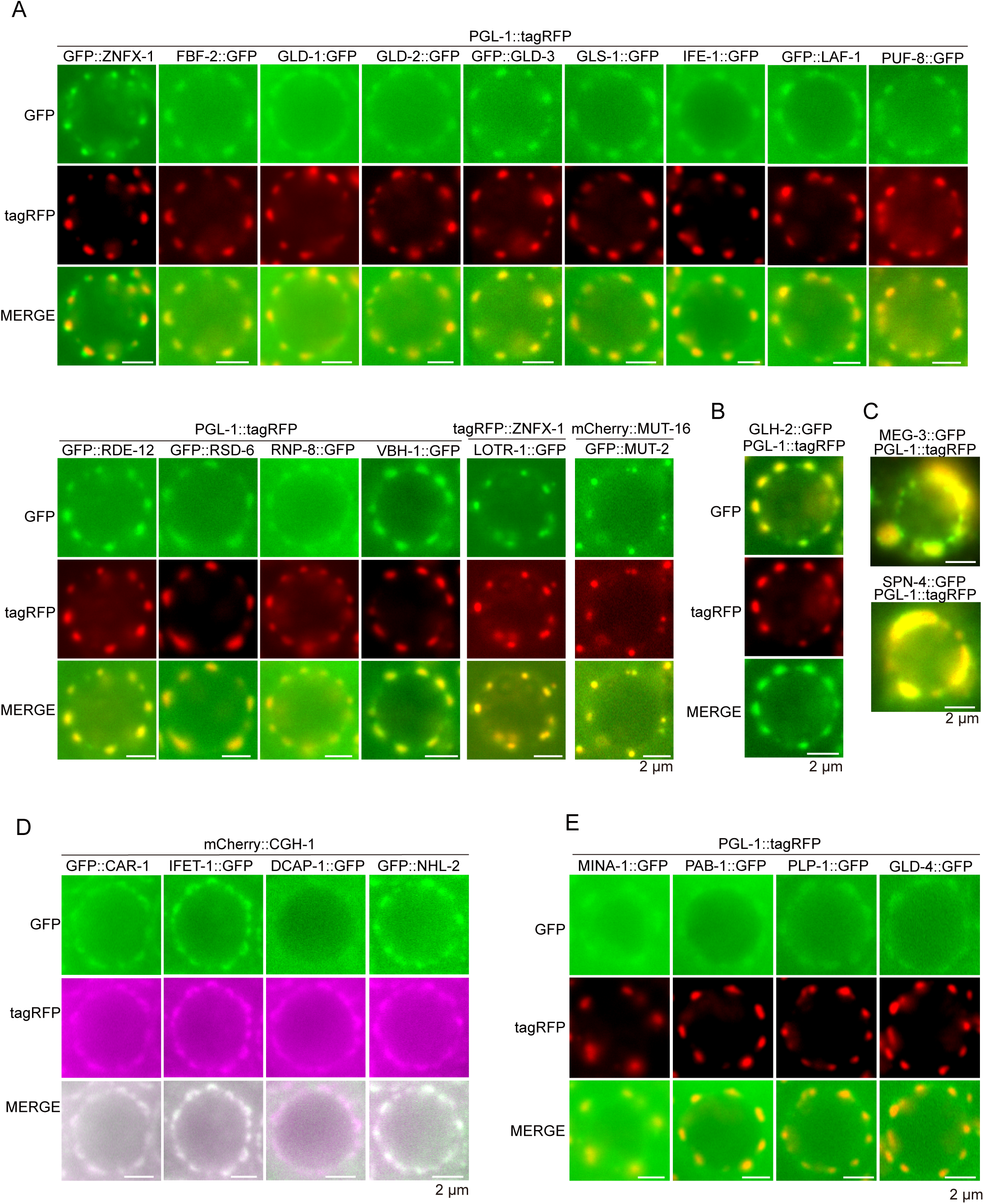
Systematic analysis of the subcellular localization of perinuclear proteins. (A, B) Fluorescence micrographs of pachytene germ cells from animals expressing the indicated proteins. (C) Fluorescence micrographs of embryo germ cells from animals expressing MEG-3::GFP and PGL-1::tagRFP, or SPN-4::tagRFP and PGL-1::tagRFP. MEG-3 and SPN-4 were exclusively expressed during embryogenesis. (D) Fluorescence micrographs of pachytene germ cells from animals expressing the indicated proteins. Perinuclear CGH-1, CAR-1, IFET-1 and NHL-2 were enriched in the P-body in germ cells. (E) Images of MINA-1::GFP, PAB-1::GFP, PLP-1::GFP GLD-4::GFP and the P-granule marker PGL-1::tagRFP in germ cells.

**Figure S3.**
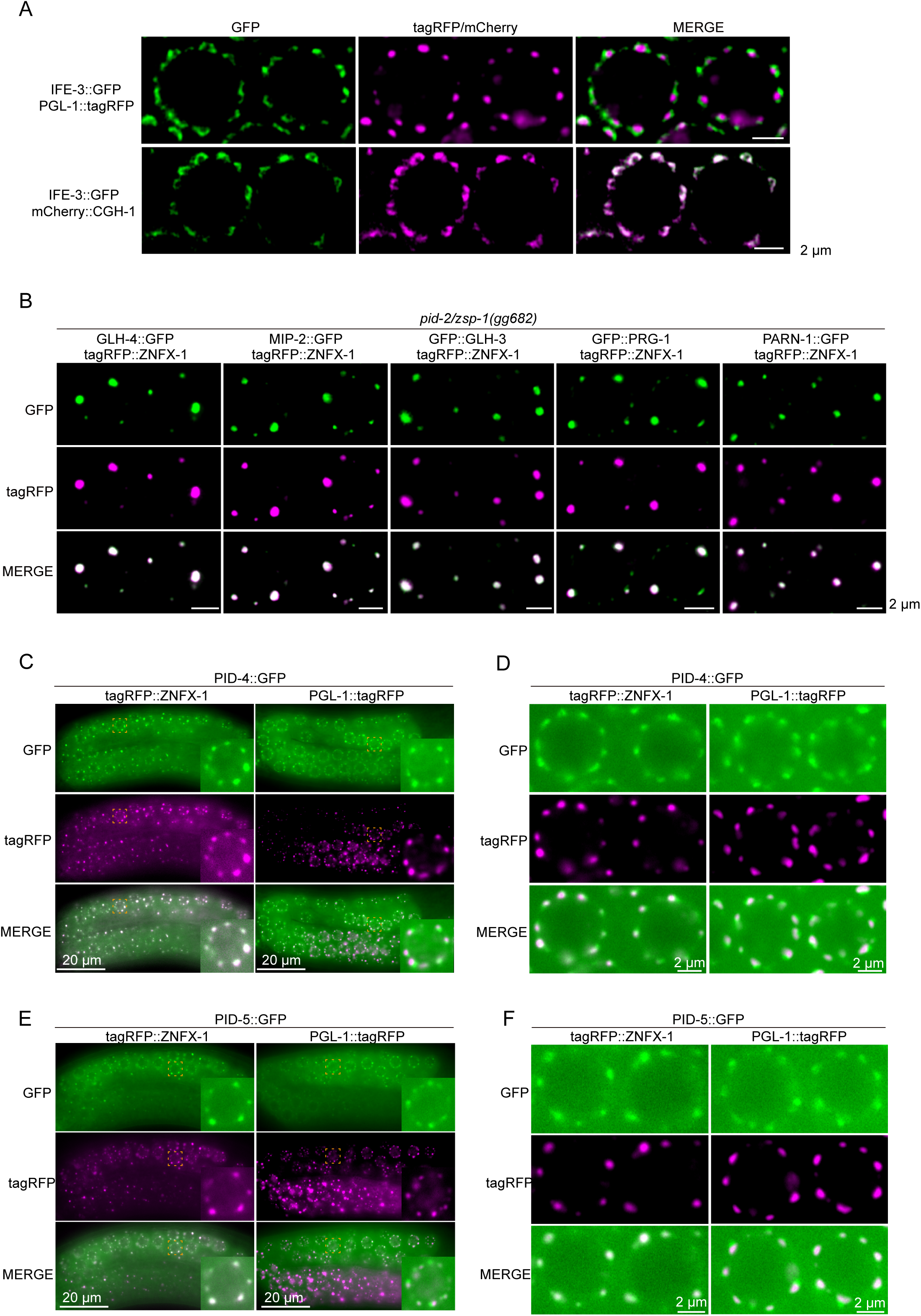
PARN-1, PRG-1, PID-4 and PID-5 were enriched in the Z granule. (A) Fluorescent images of animals expressing IFE-3::GFP and PGL-1::tagRFP, or IFE-3::GFP and mCherry::CGH-1. IFE-3 was mainly enriched in the P-body. (B) Images of tagRFP::ZNFX-1 and the indicated GFP-tagged proteins in the germ cells of *pid-2/zsp-1*(gg682) animals. PID-2 is exclusively required for Z granule homeostasis, the depletion of which results in the formation of unevenly sized Z granules around the nucleus^2,3^. GLH-3, GLH-4, MIP-2, PARN-1 and PRG-1 remained localized in the Z granules of the *mip-1* mutants. (C, D) Fluorescence micrographs of larval (C) and adult (D) animals expressing PID-4::GFP and tagRFP::ZNFX-1. PID-4 is enriched in the Z granule. (E, F) Fluorescence micrographs of larval (E) and adult (F) animals expressing PID-5::GFP and tagRFP::ZNFX-1. PID-5 is enriched in the Z granule.

**Figure S4.**
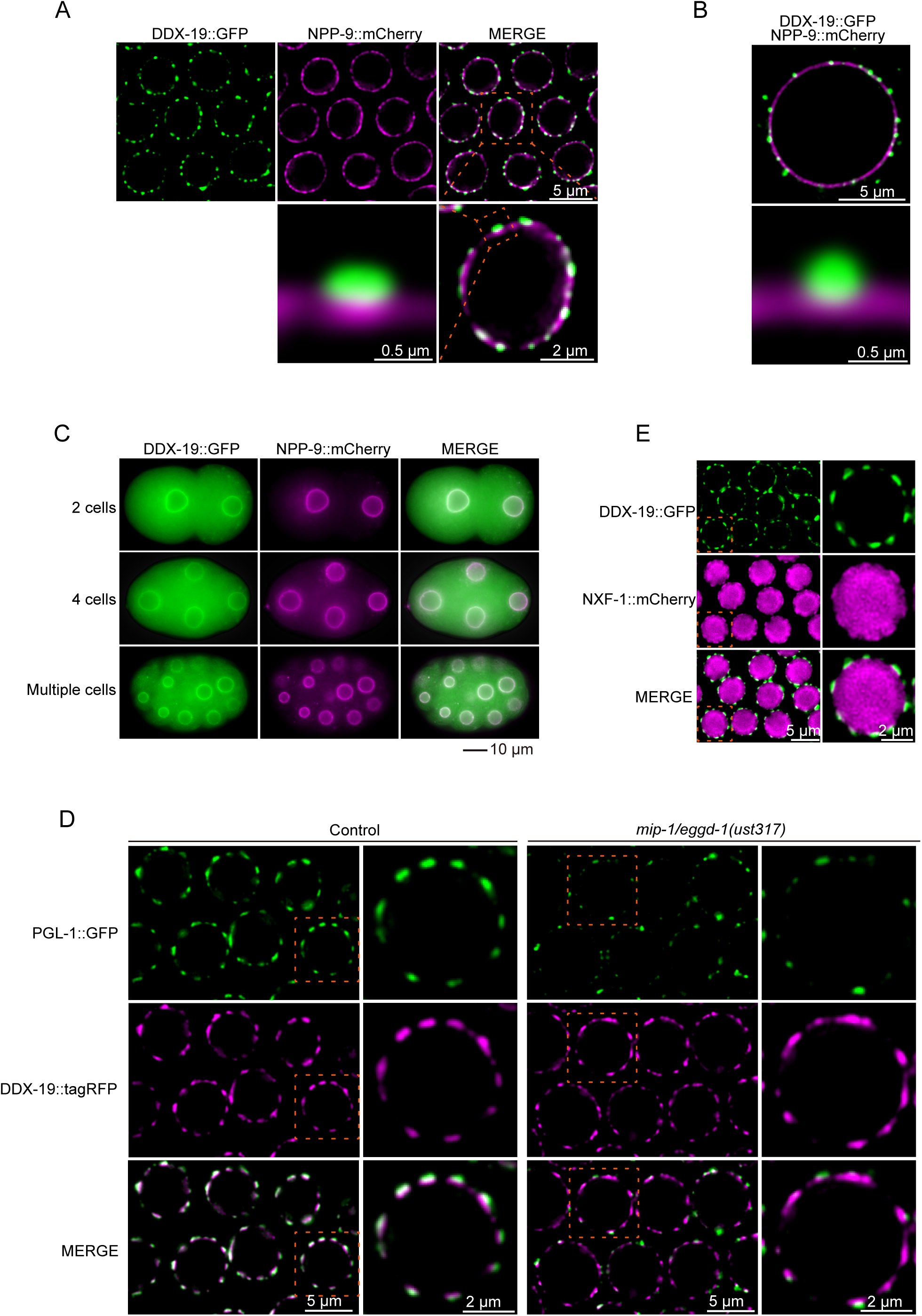
DDX-19 localized to a distinct germ granule compartment. (A, B) Fluorescent images of pachytene-stage germ cells (A) and oocytes (B) of animals expressing DDX-19::GFP and NPP-9::mCherry. DDX-19 does not colocalize with NPP-9 and is mainly enriched in the perinuclear condensates in germ cells. (C) Fluorescence micrographs of embryos expressing DDX-19::GFP and NPP-9::mCherry. DDX-19 colocalized with NPP-9 during early embryo development. (D) Fluorescent micrographs of pachytene germ cells from animals expressing DDX-19::mCherry and PGL-1::GFP in the indicated animals. (E) Images of pachytene germ cells expressing DDX-19::GFP and NXF-1::mCherry. Perinuclear NXF-1 does not colocalize with DDX-19 and may be enrich in NPCs^4^.

**Figure S5.**
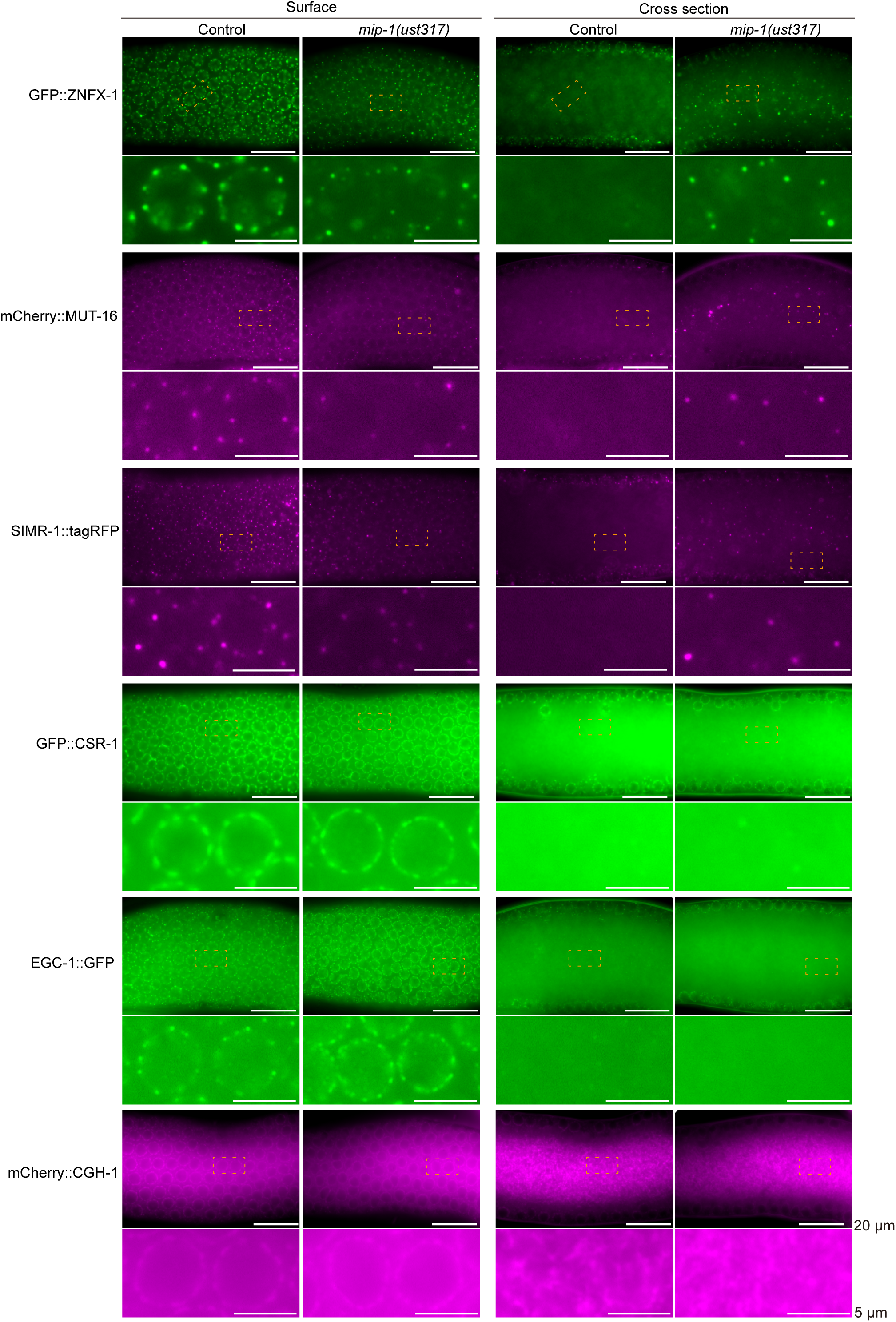
The loss of MIP-1 did not affect the accumulation of CSR-1, EGC-1 or CGH-1 in perinuclear granules. Fluorescence micrographs of wild-type and *mip-1(-)* animals expressing the indicated proteins on the surface and in rachis of adult germlines.

**Table S1.** The library of genetically modified nematode strains expressing peri-nuclear proteins labeled with fluorescent proteins.

| Gene | Tag | Loci | Allele | Strain |
| --- | --- | --- | --- | --- |
| *car-1* | *3xflag::gfp* | 5' in situ | *car-1(ust408[3xflag::gfp::car-1]) I* | SHG2239 |
| *cgh-1* | *gfp* | 5' in situ | *cgh-1(ustIS217[gfp::cgh-1]) III* | SHG1686 |
| *csr-1* | *3xflag::gfp* | 5' in situ | *csr-1(ust371[3xflag::gfp::csr-1]) IV* | SHG2040 |
| *csr-1* | *tagRFP* | 5' in situ | *csr-1(ust583[tagRFP::csr-1]) IV* | SHG2900 |
| *csr-1* | *mCherry* | 5' in situ | *csr-1(ust400[mCherry::csr-1]) IV* | SHG2901 |
| *dcap-1* | *gfp::3xflag* | 3' in situ | *dcap-1(ust409[dcap-1::gfp::3xflag]) IV* | SHG2240 |
| *ddx-19* | *gfp::3xflag* | 3' in situ | *ddx-19(ust399[ddx-19::gfp::3xflag]) II* | SHG2207 |
| *ddx-19* | *tagRFP* | 3' in situ | *ddx-19(ust584[ddx-19::tagRFP]) II* | SHG2902 |
| *ddx-19* | *mCherry* | 3' in situ | *ddx-19(ust374[ddx-19::mCherry]) II* | SHG2202 |
| *deps-1* | *gfp::3xflag* | 3' in situ | *deps-1(ust496[deps-1::gfp::3xflag]) I* | SHG2538 |
| *drh-3* | *gfp::3xflag* | 3' in situ | *drh-3(ust352[drh-3::gfp::3xflag]) I* | SHG1676 |
| *edc-3* | *3xflag::mCherry* | 5' in situ | *edc-3(ustIS218[3xflag::mCherry::edc-3]) I* | SHG1687 |
| *egc-1* | *gfp::3xflag* | 3' in situ | *egc-1(ust355[egc-1::gfp::3xflag]) III* | SHG1683 |
| *egc-1* | *mCherry* | 3' in situ | *egc-1(ust355[egc-1::mCherry]) III* | SHG1685 |
| *ego-1* | *gfp* | 5' in situ | *ego-1(ust351[gfp::ego-1]) I* | SHG1675 |
| *ego-1* | *mCherry* | 5' in situ | *ego-1(ust356[mCherry::ego-1]) I* | SHG1936 |
| *ekl-1* | *gfp::3xflag* | 3' in situ | *ekl-1(ust353[ekl-1::gfp::3xflag]) I* | SHG1679 |
| *elli-1* | *gfp::3xflag* | 3' in situ | *elli-1(ust354[elli-1::gfp::3xflag]) IV* | SHG1682 |
| *elli-1* | *tagRFP* | 5' Ex situ I | *ustIS268[mex-5p::tagrfp::elli-1::tbb-2_3'UTR] I* | SHG1941 |
| *erh-2* | *gfp::3xflag* | 3' Ex situ II | *ustIS73(erh-2p::erh-2::gfp::3xflag::erh-2_3'UTR) II* | SHG665 |
| *fbf-2* | *gfp::3xflag* | 3' in situ | *fbf-2(ust337[fbf-2::gfp::3xflag]) II* | SHG2152 |
| *gld-1* | *gfp::3xflag* | 3' in situ | *gld-1(ust451[gld-1::gfp::3xflag]) I* | SHG2362 |
| *gld-2* | *gfp::3xflag* | 3' in situ | *gld-2(ust546[gld-2::gfp::3xflag]) I* | SHG2672 |
| *gld-3* | *3xflag::gfp* | 5' in situ | *gld-3(ust495[3xflag::gfp::gld-3]) II* | SHG2537 |
| *gld-4* | *gfp::3xflag* | 3' in situ | *gld-4(ust498[gld-4::gfp::3xflag]) I* | SHG2540 |
| *gld-4* | *3xflag::gfp* | 5' in situ | *gld-4(ust593[3xflag::gfp::gld-4]) I* | SHG2957 |
| *glh-1* | *gfp::3xflag* | 5' in situ | *glh-1(ust338[3xflag::gfp::glh-1]) I* | SHG833 |
| *glh-1* | *mCherry* | 5' in situ | *glh-1(ust406[mCherry::glh-1]) I* | SHG2848 |
| *glh-2* | *gfp::3xflag* | 3' in situ | *glh-2(ust448[glh-2::gfp::3xflag]) I* | SHG2429 |
| *glh-3* | *3xflag::gfp* | 3' in situ | *glh-3(ust494[3xflag::gfp::glh-3]) I* | SHG2536 |
| *glh-4* | *gfp::3xflag* | 3' in situ | *glh-4(ust449[glh-4::gfp::3xflag]) I* | SHG2430 |
| *gls-1* | *gfp::3xflag* | 3' in situ | *gls-1(ust518[glh-1::gfp::3xflag]) I* | SHG2609 |
| *henn-1* | *gfp::3xflag* | 3' in situ | *henn-1(ust413[henn-1::gfp::3xflag]) III* | SHG2244 |
| *ife-1* | *gfp::3xflag* | 3' in situ | *ife-1(ust446[ife-1::gfp::3xflag]) III* | SHG2427 |
| *ife-3* | *gfp::3xflag* | 3' in situ | *ife-3(ustIS78[ife-3::gfp::3xflag]) V* | SHG682 |
| *ifet-1* | *gfp::3xflag* | 3' in situ | *ifet-1(ust412[ifet-1::gfp::3xflag]) III* | SHG2243 |
| *ifet-1* | *mCherry* | 3' in situ | *ifet-1(ust547[ifet-1::mcherry]) III* | SHG2766 |
| *laf-1* | *3xflag::gfp* | 5' in situ | *laf-1(ust492[3xflag::gfp::laf-1]) III* | SHG2534 |
| *lotr-1* | *gfp::3xflag* | 3' in situ | *lotr-1(ust585[lotr-1::gfp::3xflag]) I* | SHG2903 |
| *meg-3* | *gfp::3xflag* | 3' in situ | *meg-3(ust407[meg-3::gfp::3xflag]) X* | SHG2238 |
| *mina-1* | *gfp::3xflag* | 3' in situ | *mina-1(ust433[mina-1::gfp::3xflag]) I* | SHG2329 |

Table S1 Continued
| Gene | Tag | Loci | Allele | Strain |
| --- | --- | --- | --- | --- |
| *mip-2* | *gfp::3xflag* | 3' in situ | *mip-2(ust450[mip-2::gfp::3xflag]) V* | SHG2361 |
| *mut-16* | *mCherry* | 5' in situ | *mut-16(ust366[mCherry::mut-16]) I* | SHG2030 |
| *mut-2* | *3xflag::gfp* | 5' in situ | *mut-2(ust447[3xflag::gfp::mut-2]) I* | SHG2428 |
| *nhl-2* | *3xflag::gfp* | 5' in situ | *nhl-2(ust586[3xflag::gfp::nhl-2]) III* | SHG2701 |
| *npp-9* | *mCherry* | 3' in situ | *npp-9(ust373[npp-9::mCherry]) III* | SHG2190 |
| *npp-9* | *tagRFP* | 5' in situ | *npp-9(ust587[tagRFP::npp-9]) III* | SHG2904 |
| *nxf-1* | *mCherry* | 3' in situ | *nxf-1(ust372[nxf-1::mcherry]) V* | SHG2141 |
| *pab-1* | *gfp::3xflag* | 3' in situ | *pab-1(ust410[pab-1::gfp::3xflag]) I* | SHG2241 |
| *parn-1* | *gfp::3xflag* | 3' in situ | *parn-1(ust411[parn-1::gfp::3xflag]) V* | SHG2242 |
| *pgl-1* | *gfp::3xflag* | 3' in situ | *pgl-1(ust588[pgl-1::gfp::3xflag]) IV* | SHG2905 |
| *pgl-1* | *bfp* | 3' in situ | *pgl-1(ust363[pgl-1::bfp]) IV* | SHG2028 |
| *pgl-2* | *3xflag::gfp* | 3' in situ | *pgl-2(ust515[3xflag::gfp::pgl-2]) III* | SHG2606 |
| *pgl-3* | *gfp::3xflag* | 3' in situ | *pgl-3(ust437[pgl-3::gfp::3xflag]) V* | SHG2333 |
| *pics-1/pid-3* | *gfp::3xflag* | 3' Ex situ II | *ustIS60(pics-1p::pics-1::gfp::3xflag::pics-1_3'UTR) II* | SHG542 |
| *pid-1* | *gfp::3xflag* | 3' Ex situ II | *ustIS55(pid-1p::pid-1::gfp::3xflag::pid-1_3'UTR) II* | SHG670 |
| *pid-4* | *gfp::3xflag* | 3' in situ | *pid-4(ust516[pid-4::gfp::3xflag]) I* | SHG2607 |
| *pid-5* | *gfp::3xflag* | 3' in situ | *pid-5(ust517[pid-5::gfp::3xflag]) V* | SHG2608 |
| *prg-1* | *3xflag::gfp* | 5' in situ | *prg-1(ustIS81[3xflag::gfp::prg-1]) I* | SHG763 |
| *plp-1* | *gfp::3xflag* | 3' in situ | *plp-1(ust432[plp-1::gfp::3xflag]) IV* | SHG2328 |
| *puf-8* | *gfp::3xflag* | 3' in situ | *puf-8(ust245[puf-8::gfp::3xflag]) II* | SHG1614 |
| *rde-12* | *3xflag::gfp* | 5' in situ | *rde-12(ust575[3xflag::gfp::rde-12]) V* | SHG2882 |
| *rnp-8* | *gfp::3xflag* | 3' in situ | *rnp-9(ust589[rnp-8::gfp::3xflag]) I* | SHG2906 |
| *rrf-1* | *gfp::3xflag* | 3' in situ | *rrf-1(ust590[rrf-1::gfp::3xflag]) I* | SHG2907 |
| *rrf-1* | *tagRFP* | 3' in situ | *rrf-1(ust364[rrf-1::ragRFP]) I* | SHG2673 |
| *rsd-6* | *3xflag::gfp* | 5' in situ | *rsd-6(ust591[3xflag::gfp::rsd-6]) I* | SHG2908 |
| *simr-1* | *tagRFP* | 3' in situ | *simr-1(ust365[simr-1::tagRFP]) I* | SHG2440 |
| *spn-4* | *gfp::3xflag* | 3' in situ | *spn-4(ust434[spn-4::gfp::3xflag]) V* | SHG2330 |
| *tofu-6* | *gfp::3xflag* | 3' Ex situ II | *ustIS32(tofu-6p::tofu-6::gfp::3xflag::tofu-6_3'UTR) II* | SHG357 |
| *vbh-1* | *gfp::3xflag* | 3' in situ | *vbh-1(ust435[vbh-1::gfp::3xflag]) I* | SHG2331 |
| *wago-1* | *3xflag::gfp* | 5' in situ | *ustIS106(wago-1p::3xflag::gfp::wago-1::wago-1_3'UTR) II* | SHG659 |
| *wago-3* | *gfp::3xflag* | 3' Ex situ II | *ustIS64(wago-3p::3xflag::gfp::wago-3::wago-3_3'UTR) II* | SHG578 |
| *wago-4* | *3xflag::gfp* | 5' Ex situ II | *ustIS25(wago-4p::3xflag::gfp::wago-4::wago-4_3'UTR) II* | SHG487 |
| *znfx-1* | *mCherry* | 5' in situ | *znfx-1(ust362[mCherry::znfx-1]) V* | SHG2909 |

**Table S2.**
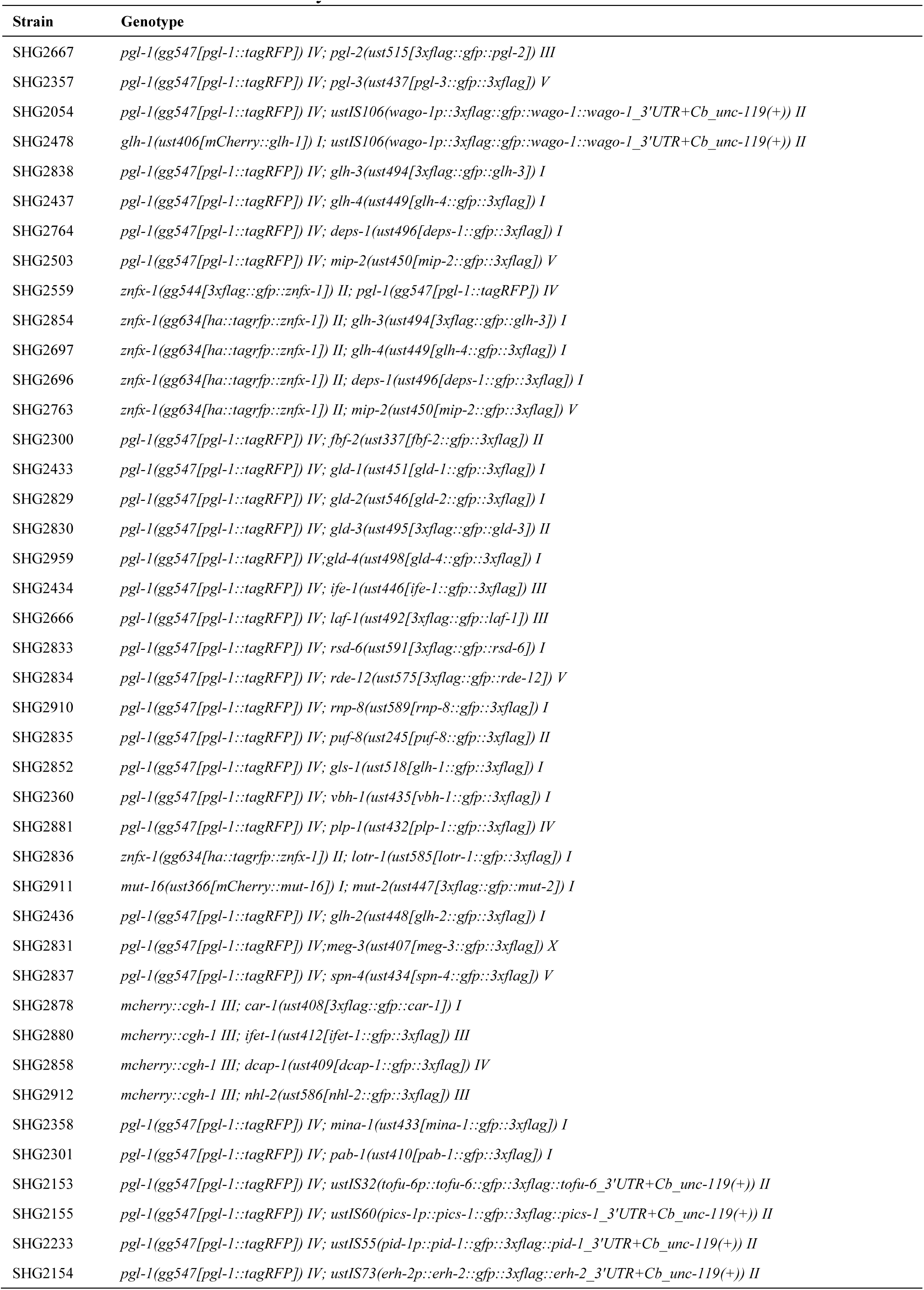
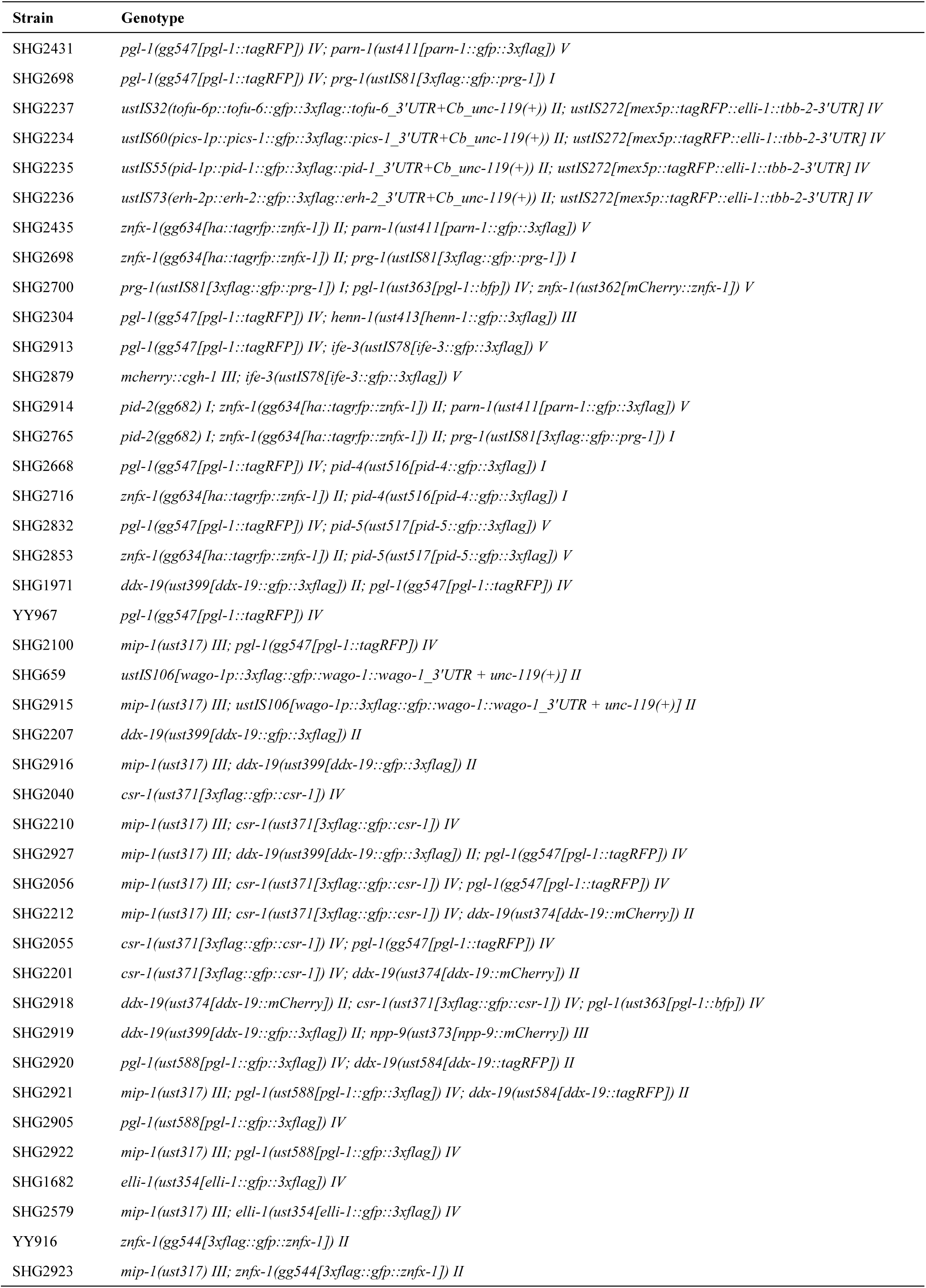
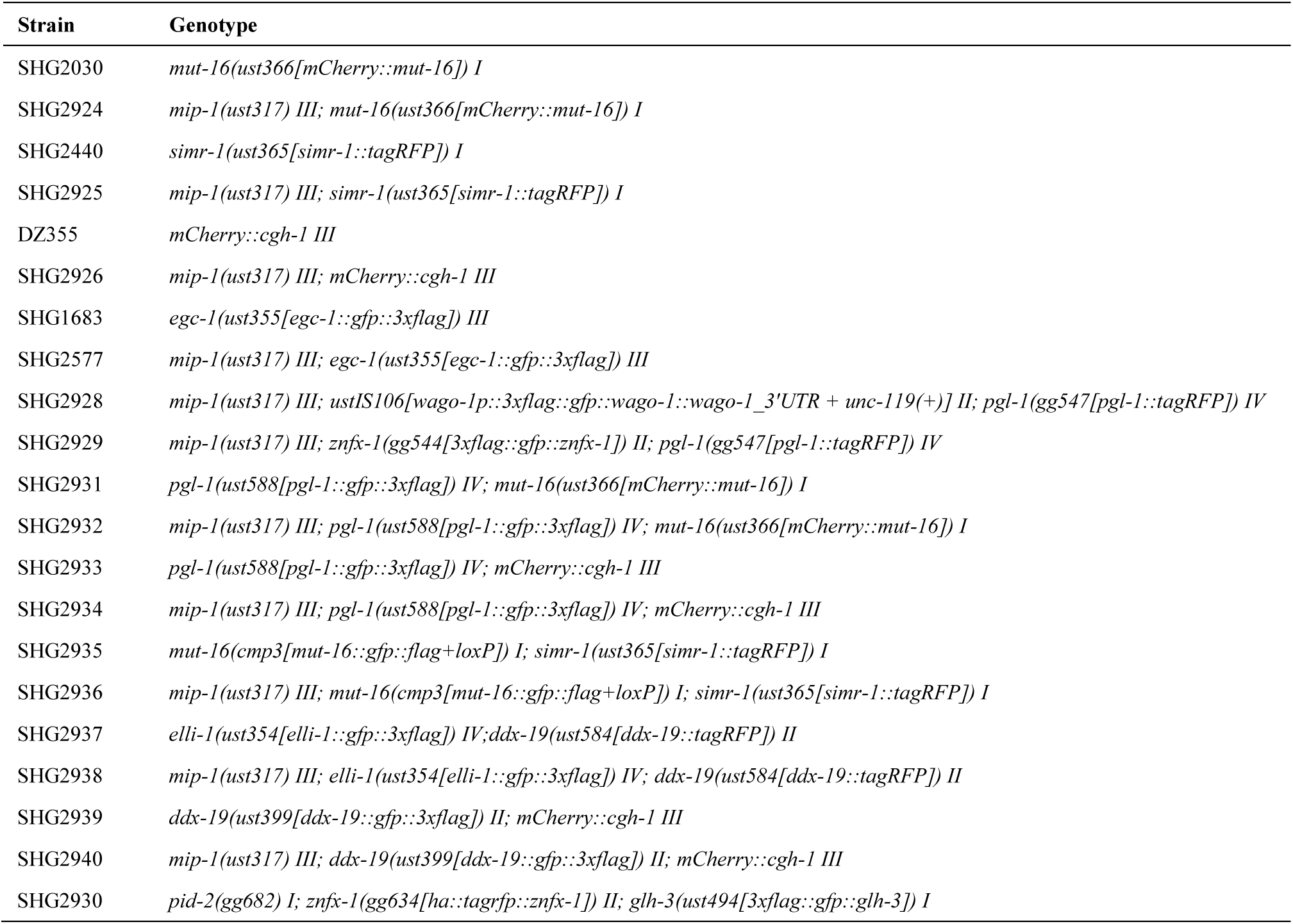
The list of strains used in this study.

## References

1. Shin, Y., and Brangwynne, C.P. (2017). Liquid phase condensation in cell physiology and disease. Science 357. 10.1126/science.aaf4382.

2. Courchaine, E.M., Lu, A., and Neugebauer, K.M. (2016). Droplet organelles? Embo Journal 35, 1603–1612. 10.15252/embj.201593517.

3. Alberti, S., and Dormann, D. (2019). Liquid-Liquid Phase Separation in Disease. Annual Review of Genetics, Vol 53 53, 171–194. 10.1146/annurev-genet-112618-043527.

4. Banani, S.F., Lee, H.O., Hyman, A.A., and Rosen, M.K. (2017). Biomolecular condensates: organizers of cellular biochemistry. Nature Reviews Molecular Cell Biology 18, 285–298. 10.1038/nrm.2017.7.

5. Toretsky, J.A., and Wright, P.E. (2014). Assemblages: Functional units formed by cellular phase separation. Journal of Cell Biology 206, 579–588. 10.1083/jcb.201404124.

6. Standart, N., and Weil, D. (2018). P-Bodies: Cytosolic Droplets for Coordinated mRNA Storage. Trends in Genetics 34, 612–626. 10.1016/j.tig.2018.05.005.

7. Fare, C.M., Villani, A., Drake, L.E., and Shorter, J. (2021). Higher-order organization of biomolecular condensates. Open Biology 11. 10.1098/rsob.210137.

8. Lafontaine, D.L.J., Riback, J.A., Bascetin, R., and Brangwynne, C.P. (2021). The nucleolus as a multiphase liquid condensate. Nature Reviews Molecular Cell Biology 22, 165–182. 10.1038/s41580-020-0272-6.

9. Yanagawa, M., and Shimobayashi, S.F. (2023). Multi-dimensional condensation of intracellular biomolecules. Journal of Biochemistry 175, 179–186. 10.1093/jb/mvad095.

10. Feric, M., Vaidya, N., Harmon, T.S., Mitrea, D.M., Zhu, L., Richardson, T.M., Kriwacki, R.W., Pappu, R.V., and Brangwynne, C.P. (2016). Coexisting Liquid Phases Underlie Nucleolar Subcompartments. Cell 165, 1686–1697. 10.1016/j.cell.2016.04.047.

11. West, J.A., Mito, M., Kurosaka, S., Takumi, T., Tanegashima, C., Chujo, T., Yanaka, K., Kingston, R.E., Hirose, T., Bond, C., et al. (2016). Structural, super-resolution microscopy analysis of paraspeckle nuclear body organization. Journal of Cell Biology 214, 817–830. 10.1083/jcb.201601071.

12. Fei, J.Y., Jadaliha, M., Harmon, T.S., Li, I.T.S., Hua, B.Y., Hao, Q.Y., Holehouse, A.S., Reyer, M., Sun, Q.Y., Freier, S.M., et al. (2017). Quantitative analysis of multilayer organization of proteins and RNA in nuclear speckles at super resolution. Journal of Cell Science 130, 4180–4192. 10.1242/jcs.206854.

13. Jain, S., Wheeler, J.R., Walters, R.W., Agrawal, A., Barsic, A., and Parker, R. (2016). ATPase-Modulated Stress Granules Contain a Diverse Proteome and Substructure. Cell 164, 487–498. 10.1016/j.cell.2015.12.038.

14. Wan, G., Fields, B.D., Spracklin, G., Shukla, A., Phillips, C.M., and Kennedy, S. (2018). Spatiotemporal regulation of liquid-like condensates in epigenetic inheritance. Nature 557, 679–683. 10.1038/s41586-018-0132-0.

15. Wan, G., Bajaj, L., Fields, B., Dodson, A.E., Pagano, D., Fei, Y.H., and Kennedy, S. (2021). ZSP-1 is a Z granule surface protein required for Z granule fluidity and germline immortality in *Caenorhabditis elegans*. Embo Journal 40. 10.15252/embj.2020105612.

16. Shan, L., Xu, G., Yao, R.W., Luan, P.F., Huang, Y.K., Zhang, P.H., Pan, Y.H., Zhang, L., Gao, X., Li, Y., et al. (2023). Nucleolar URB1 ensures 3′ ETS rRNA removal to prevent exosome surveillance. Nature 615, 526–534. 10.1038/s41586-023-05767-5.

17. Eddy, E.M. (1974). Fine structural observations on the form and distribution of nuage in germ cells of the rat. Anat Rec 178, 731–757. 10.1002/ar.1091780406.

18. Eddy, E.M. (1975). Germ plasm and the differentiation of the germ cell line. Int Rev Cytol 43, 229–280. 10.1016/s0074-7696(08)60070-4.

19. Voronina, E., Seydoux, G., Sassone-Corsi, P., and Nagamori, I. (2011). RNA Granules in Germ Cells. Cold Spring Harbor Perspectives in Biology 3. 10.1101/cshperspect.a002774.

20. Trcek, T., and Lehmann, R. (2019). Germ granules in *Drosophila*. Traffic 20, 650–660. 10.1111/tra.12674.

21. Phillips, C.M., and Updike, D.L. (2022). Germ granules and gene regulation in the germline. Genetics 220. 10.1093/genetics/iyab195.

22. Kemph, A., and Lynch, J.A. (2022). Evolution of germ plasm assembly and function among the insects. Current Opinion in Insect Science 50. 10.1016/j.cois.2022.100883.

23. Dodson, A.E., and Kennedy, S. (2020). Phase Separation in Germ Cells and Development. Developmental Cell 55, 4–17. 10.1016/j.devcel.2020.09.004.

24. Lehmann, R. (2016). Germ Plasm Biogenesis-An Oskar-Centric Perspective. Essays on Developmental Biology, Pt A 116, 679–707. 10.1016/bs.ctdb.2015.11.024.

25. Mukherjee, N., and Mukherjee, C. (2021). Germ cell ribonucleoprotein granules in different clades of life: From insects to mammals. Wiley Interdisciplinary Reviews-Rna 12. 10.1002/wrna.1642.

26. Du, Z., Shi, K., Brown, J.S., He, T., Wu, W.S., Zhang, Y., Lee, H.C., and Zhang, D.L. (2023). Condensate cooperativity underlies transgenerational gene silencing. Cell Reports 42. 10.1016/j.celrep.2023.112859.

27. Chen., X., Wang., K., Mufti., F.U.D., Xu., D., Zhu., C., Huang., X., Zeng., C., Jin., Q., Huang., X., Yan., Y.-h., et al. (2023). Germ granule compartments coordinate specialized small RNA production. bioRxiv. 10.1101/2023.12.04.570003.

28. Brangwynne, C.P., Eckmann, C.R., Courson, D.S., Rybarska, A., Hoege, C., Gharakhani, J., Jülicher, F., and Hyman, A.A. (2009). Germline P Granules Are Liquid Droplets That Localize by Controlled Dissolution/Condensation. Science 324, 1729–1732. 10.1126/science.1172046.

29. Seydoux, G. (2018). The P Granules of C. elegans: A Genetic Model for the Study of RNA-Protein Condensates. Journal of Molecular Biology 430, 4702–4710. 10.1016/j.jmb.2018.08.007.

30. Updike, D., and Strome, S. (2010). P granule assembly and function in *Caenorhabditis elegans* germ cells. Journal of Andrology 31, 53–60. 10.2164/jandrol.109.008292.

31. Xu, F., Feng, X.Z., Chen, X.Y., Weng, C.C., Yan, Q., Xu, T., Hong, M.J., and Guang, S.H. (2018). A Cytoplasmic Argonaute Protein Promotes the Inheritance of RNAi. Cell Reports 23, 2482–2494. 10.1016/j.celrep.2018.04.072.

32. Ouyang, J.P.T., Zhang, W.L., and Seydoux, G. (2022). The conserved helicase ZNFX-1 memorializes silenced RNAs in perinuclear condensates. Nature Cell Biology 24, 1129–1140. 10.1038/s41556-022-00940-w.

33. Zhang, C., Montgomery, T.A., Gabel, H.W., Fischer, S.E.J., Phillips, C.M., Fahlgren, N., Sullivan, C.M., Carrington, J.C., and Ruvkun, G. (2011). mut-16 and other mutator class genes modulate 22G and 26G siRNA pathways in *Caenorhabditis elegans*. Proceedings of the National Academy of Sciences of the United States of America 108, 1201–1208. 10.1073/pnas.1018695108.

34. Phillips, C.M., Montgomery, T.A., Breen, P.C., and Ruvkun, G. (2012). MUT-16 promotes formation of perinuclear mutator foci required for RNA silencing in the *C. elegans* germline. Genes & Development 26, 1433–1444. 10.1101/gad.193904.112.

35. Manage, K.I., Rogers, A.K., Wallis, D.C., Liebel, C.J., Anderson, D.C., Nguyen, D.A.H., Arca, K., Brown, K.C., Rodrigues, R.J.C., de Albuquerque, B.F.M., et al. (2020). A tudor domain protein, SIMR-1, promotes siRNA production at piRNA-targeted mRNAs in. Elife 9. 10.7554/eLife.56731.

36. Chen, S.H., and Phillips, C.M. (2024). HRDE-2 drives small RNA specificity for the nuclear Argonaute protein HRDE-1. Nature Communications 15. 10.1038/s41467-024-45245-8.

37. Gallo, C.M., Munro, E., Rasoloson, D., Merritt, C., and Seydoux, G. (2008). Processing bodies and germ granules are distinct RNA granules that interact in *C. elegans* embryos. Developmental Biology 323, 76–87. 10.1016/j.ydbio.2008.07.008.

38. Cassani, M., and Seydoux, G. (2024). P-body-like condensates in the germline. Seminars in Cell & Developmental Biology 157, 24–32. 10.1016/j.semcdb.2023.06.010.

39. Uebel, C.J., Rajeev, S., and Phillips, C.M. (2023). *Caenorhabditis elegans* germ granules are present in distinct configurations and assemble in a hierarchical manner. Development 150. 10.1242/dev.202284.

40. Sundby, A.E., Molnar, R.I., and Claycomb, J.M. (2021). Connecting the Dots: Linking Small RNA Pathways and Germ Granules. Trends in Cell Biology 31, 387–401. 10.1016/j.tcb.2020.12.012.

41. Uebel, C.J., Anderson, D.C., Mandarino, L.M., Manage, K.I., Aynaszyan, S., and Phillips, C.M. (2018). Distinct regions of the intrinsically disordered protein MUT-16 mediate assembly of a small RNA amplification complex and promote phase separation of Mutator foci. Plos Genetics 14. 10.1371/journal.pgen.1007542.

42. Shukla, A., Yen, J., Pagano, D.J., Dodso, A.E., Fei, Y.H., Gorham, J., Seidman, J.G., Wickens, M., and Kennedy, S. (2020). poly(UG)-tailed RNAs in genome protection and epigenetic inheritance. Nature 582, 283–288. 10.1038/s41586-020-2323-8.

43. Placentino, M., Domingues, A.M.D., Schreier, J., Dietz, S., Hellmann, S., de Albuquerque, B.F.M., Butter, F., and Ketting, R.F. (2021). Intrinsically disordered protein PID-2 modulates Z granules and is required for heritable piRNA-induced silencing in the *Caenorhabditis elegans* embryo. Embo Journal 40. 10.15252/embj.2020105280.

44. Marnik, E.A., Almeida, M.V., Cipriani, P.G., Chung, G.R., Caspani, E., Karaulanov, E., Gan, H.H., Zinno, J., Isolehto, I.J., Kielisch, F., et al. (2022). The Caenorhabditis elegans TDRD5/7-like protein, LOTR-1, interacts with the helicase ZNFX-1 to balance epigenetic signals in the germline. Plos Genetics 18. 10.1371/journal.pgen.1010245.

45. Cipriani, P.G., Bay, O., Zinno, J., Gutwein, M., Gan, H.H., Mayya, V.K., Chung, G., Chen, J.X., Fahs, H., Guan, Y., et al. (2021). Novel LOTUS-domain proteins are organizational hubs that recruit *C. elegans* Vasa to germ granules. Elife 10. 10.7554/eLife.60833.

46. Price, I.F., Hertz, H.L., Pastore, B., Wagner, J., and Tang, W. (2021). Proximity labeling identifies LOTUS domain proteins that promote the formation of perinuclear germ granules in *C. elegans*. Elife 10. 10.7554/eLife.72276.

47. Sheth, U., Pitt, J., Dennis, S., and Priess, J.R. (2010). Perinuclear P granules are the principal sites of mRNA export in adult *C. elegans* germ cells. Development 137, 1305–1314. 10.1242/dev.044255.

48. Zeng, C.M., Weng, C.C., Wang, X.Y., Yan, Y.H., Li, W.J., Xu, D.M., Hong, M.J., Liao, S.H., Dong, M.Q., Feng, X.Z., et al. (2019). Functional Proteomics Identifies a PICS Complex Required for piRNA Maturation and Chromosome Segregation. Cell Reports 27, 3561–3572. 10.1016/j.celrep.2019.05.076.

49. Davis, G.M., Tu, S.K., Anderson, J.W.T., Colson, R.N., Gunzburg, M.J., Francisco, M.A., Ray, D., Shrubsole, S.P., Sobotka, J.A., Seroussi, U., et al. (2018). The TRIM-NHL protein NHL-2 is a co-factor in the nuclear and somatic RNAi pathways in *C. elegans*. Elife 7. 10.7554/eLife.35478.

50. Kawasaki, I., Shim, Y.H., Kirchner, J., Kaminker, J., Wood, W.B., and Strome, S. (1998). PGL-1, a predicted RNA-binding component of germ granules, is essential for fertility in *C. elegans*. Cell 94, 635–645. 10.1016/S0092-8674(00)81605-0.

51. Kawasaki, L., Amiri, A., Fan, Y., Meyer, N., Dunkelbarger, S., Motohashi, T., Karashima, T., Bossinger, O., and Strome, S. (2004). The PGL family proteins associate with germ granules and function redundantly in germline development. Genetics 167, 645–661. 10.1534/genetics.103.023093.

52. Gu, W.F., Shirayama, M., Conte, D., Vasale, J., Batista, P.J., Claycomb, J.M., Moresco, J.J., Youngman, E.M., Keys, J., Stoltz, M.J., et al. (2009). Distinct argonaute-mediated 22G-RNA pathways direct genome surveillance in the *C. elegans* germline. Molecular Cell 36, 231–244. 10.1016/j.molcel.2009.09.020.

53. Gustafson, E.A., and Wessel, G.M. (2010). Vasa genes: Emerging roles in the germ line and in multipotent cells. Bioessays 32, 626–637. 10.1002/bies.201000001.

54. Spike, C., Meyer, N., Racen, E., Orsborn, A., Kirchner, J., Kuznicki, K., Yee, C., Bennettt, K., and Strome, S. (2008). Genetic analysis of the GLH family of p-granule proteins. Genetics 178, 1973–1987. 10.1534/genetics.107.083469.

55. Chen, W.J., Brown, J.S., He, T., Wu, W.S., Tu, S.K., Weng, Z.P., Zhang, D.L., and Lee, H.C. (2022). GLH/VASA helicases promote germ granule formation to ensure the fidelity of piRNA-mediated transcriptome surveillance. Nature Communications 13. 10.1038/s41467-022-32880-2.

56. Dai, S.Y., Tang, X.Y., Li, L.L., Ishidate, T., Ozturk, A.R., Chen, H., Dube, A.L., Yan, Y.H., Dong, M.Q., Shen, E.Z., and Mello, C.C. (2022). A family of *C. elegans* VASA homologs control Argonaute pathway specificity and promote transgenerational silencing. Cell Reports 40. 10.1016/j.celrep.2022.111265.

57. Spike, C.A., Bader, J., Reinke, V., and Strome, S. (2008). DEPS-1 promotes P-granule assembly and RNA interference in *C. elegans* germ cells. Development 135, 983–993. 10.1242/dev.015552.

58. Suen, K.M., Braukmann, F., Butler, R., Bensaddek, D., Akay, A., Lin, C.C., Milonaityte, D., Doshi, N., Sapetschnig, A., Lamond, A., et al. (2020). DEPS-1 is required for piRNA-dependent silencing and PIWI condensate organisation in *Caenorhabditis elegans*. Nature Communications 11. 10.1038/s41467-020-18089-1.

59. Ketting, R.F., and Cochella, L. (2021). Concepts and functions of small RNA pathways in *C. elegans*. Nematode Models of Development and Disease 144, 45–89. 10.1016/bs.ctdb.2020.08.002.

60. Weng, C.C., Kosalka, J., Berkyurek, A.C., Stempor, P., Feng, X.Z., Mao, H., Zeng, C.M., Li, W.J., Yan, Y.H., Dong, M.Q., et al. (2019). The USTC co-opts an ancient machinery to drive piRNA transcription in *C. elegans*. Genes & Development 33, 90–102. 10.1101/gad.319293.118.

61. Podvalnaya, N., Bronkhorst, A.W., Lichtenberger, R., Hellmann, S., Nischwitz, E., Falk, T., Karaulanov, E., Butter, F., Falk, S., and Ketting, R.F. (2023). piRNA processing by a trimeric Schlafen-domain nuclease. Nature 622, 402–409. 10.1038/s41586-023-06588-2.

62. Rodrigues, R.J.C., Domingues, A.M.D., Hellmann, S., Dietz, S., de Albuquerque, B.F.M., Renz, C., Ulrich, H.D., Sarkies, P., Butter, F., and Ketting, R.F. (2019). PETISCO is a novel protein complex required for 21U RNA biogenesis and embryonic viability. Genes & Development 33, 857–870. 10.1101/gad.322446.118.

63. Tang, W., Tu, S., Lee, H.C., Weng, Z.P., and Mello, C.C. (2016). The RNase PARN-1 Trims piRNA 3’ Ends to Promote Transcriptome Surveillance in *C. elegans*. Cell 164, 974–984. 10.1016/j.cell.2016.02.008.

64. Batista, P.J., Ruby, J.G., Claycomb, J.M., Chiang, R., Fahlgren, N., Kasschau, K.D., Chaves, D.A., Gu, W.F., Vasale, J.J., Duan, S.H., et al. (2008). PRG-1 and 21U-RNAs interact to form the piRNA complex required for fertility in *C. elegans*. Molecular Cell 31, 67–78. 10.1016/j.molcel.2008.06.002.

65. Das, P.P., Bagijn, M.P., Goldstein, L.D., Woolford, J.R., Lehrbach, N.J., Sapetschnig, A., Buhecha, H.R., Gilchrist, M.J., Howe, K.L., Stark, R., et al. (2008). Piwi and piRNAs act upstream of an endogenous siRNA pathway to suppress Tc3 transposon mobility in the *Caenorhabditis elegans* germline. Molecular Cell 31, 79–90. 10.1016/j.molcel.2008.06.003.

66. Wang, G., and Reinke, V. (2008). A *C. elegans* Piwi, PRG-1, regulates 21U-RNAs during spermatogenesis. Current Biology 18, 861–867. 10.1016/j.cub.2008.05.009.

67. Stower, H. (2012). Small RNAs: piRNA surveillance in the *C. elegans* germline. Nature Reviews Genetics 13. 10.1038/nrg3289.

68. Billi, A.C., Alessi, A.F., Khivansara, V., Han, T., Freeberg, M., Mitani, S., and Kim, J.K. (2012). The HEN1 Ortholog, HENN-1, Methylates and Stabilizes Select Subclasses of Germline Small RNAs. Plos Genetics 8, 84–99. 10.1371/journal.pgen.1002617.

69. Kamminga, L.M., van Wolfswinkel, J.C., Luteijn, M.J., Kaaij, L.J.T., Bagijn, M.P., Sapetschnig, A., Miska, E.A., Berezikov, E., and Ketting, R.F. (2012). Differential impact of the HEN1 homolog HENN-1 on 21U and 26G RNAs in the germline of *Caenorhabditis elegans*. Plos Genetics 8. 10.1371/journal.pgen.1002702.

70. Montgomery, T.A., Rim, Y.S., Zhang, C., Dowen, R.H., Phillips, C.M., Fischer, S.E.J., and Ruvkun, G. (2012). PIWI associated siRNAs and piRNAs specifically require the *Caenorhabditis elegans* HEN1 ortholog henn-1. Plos Genetics 8, 72–83. 10.1371/journal.pgen.1002616.

71. Price, I.F., Wagner, J.A., Pastore, B., Hertz, H.L., and Tang, W. (2023). *C. elegans* germ granules sculpt both germline and somatic RNAome. Nature Communications 14. 10.1038/s41467-023-41556-4.

72. Charlesworth, A.G., Seroussi, U., Lehrbach, N.J., Renaud, M.S., Sundby, A.E., Molnar, R., Lao, R.X., Willis, A.R., Woock, J.R., Aber, M.J., et al. (2021). Two isoforms of the essential Argonaute CSR-1 differentially regulate sperm and oocyte fertility. Nucleic Acids Research 49, 8836–8865. 10.1093/nar/gkab619.

73. Nguyen, D.A.H., and Phillips, C.M. (2021). Arginine methylation promotes siRNA-binding specificity for a spermatogenesis-specific isoform of the Argonaute protein CSR-1. Nature Communications 12. 10.1038/s41467-021-24526-6.

74. Gerson-Gurwitz, A., Wang, S.H., Sathe, S., Green, R., Yeo, G.W., Oegema, K., and Desai, A. (2016). A Small RNA-Catalytic Argonaute Pathway Tunes Germline Transcript Levels to Ensure Embryonic Divisions. Cell 165, 396–409. 10.1016/j.cell.2016.02.040.

75. Tan, W., Zolotukhin, A.S., Bear, J., Patenaude, D.J., and Felber, B.K. (2000). The mRNA export in *Caenorhabditis elegans* is mediated by Ce-NXF-1, an ortholog of human TAP/NXF and Saccharomyces cerevisiae Mex67p. Rna 6, 1762–1772. 10.1017/S1355838200000832.

76. Zheleva, A., Gómez-Orte, E., Sáenz-Narciso, B., Ezcurra, B., Kassahun, H., de Toro, M., Miranda-Vizuete, A., Schnabel, R., Nilsen, H., and Cabello, J. (2019). Reduction of mRNA export unmasks different tissue sensitivities to low mRNA levels during *Caenorhabditis elegans* development. Plos Genetics 15. 10.1371/journal.pgen.1008338.

77. Khong, A., Matheny, T., Jain, S., Mitchell, S.F., Wheeler, J.R., and Parker, R. (2017). The Stress Granule Transcriptome Reveals Principles of mRNA Accumulation in Stress Granules. Molecular Cell 68, 808–820. 10.1016/j.molcel.2017.10.015.

78. Markmiller, S., Soltanieh, S., Server, K.L., Mak, R., Jin, W.H., Fang, M.Y., Luo, E.C., Krach, F., Yang, D.J., Sen, A., et al. (2018). Context-Dependent and Disease-Specific Diversity in Protein Interactions within Stress Granules. Cell 172, 590–604. 10.1016/j.cell.2017.12.032.

79. Youn, J.Y., Dunham, W.H., Hong, S.J., Knight, J.D.R., Bashkurov, M., Chen, G.I., Bagci, H., Rathod, B., MacLeod, G., Eng, S.W.M., et al. (2018). High-Density Proximity Mapping Reveals the Subcellular Organization of mRNA-Associated Granules and Bodies. Molecular Cell 69, 517–532. 10.1016/j.molcel.2017.12.020.

80. Updike, D.L., Hachey, S.J., Kreher, J., and Strome, S. (2011). P granules extend the nuclear pore complex environment in the *C. elegans* germ line. Journal of Cell Biology 192, 939–948. 10.1083/jcb.201010104.

81. Pastore, B., Hertz, H.L., Price, I.F., and Tang, W. (2021). pre-piRNA trimming and 2′-O-methylation protect piRNAs from 3′ tailing and degradation in *C. elegans*. Cell Reports 36. 10.1016/j.celrep.2021.109640.

82. Tanaka, T., Hosokawa, M., Vagin, V.V., Reuter, M., Hayashi, E., Mochizuki, A.L., Kitamura, K., Yamanaka, H., Kondoh, G., Okawa, K., et al. (2011). Tudor domain containing 7 (Tdrd7) is essential for dynamic ribonucleoprotein (RNP) remodeling of chromatoid bodies during spermatogenesis. Proceedings of the National Academy of Sciences of the United States of America 108, 10579–10584. 10.1073/pnas.1015447108.

83. Cohen-Fix, O., and Askjaer, P. (2017). Cell Biology of the *Caenorhabditis elegans* Nucleus. Genetics 205, 25–59. 10.1534/genetics.116.197160.

84. Pitt, J.N., Schisa, J.A., and Priess, J.R. (2000). P Granules in the germ cells of *Caenorhabditis elegans* adults are associated with clusters of nuclear pores and contain RNA. Developmental Biology 219, 315–333. 10.1006/dbio.2000.9607.

85. Snay-Hodge, C.A., Colot, H.V., Goldstein, A.L., and Cole, C.N. (1998). Dbp5p/Rat8p is a yeast nuclear pore-associated DEAD-box protein essential for RNA export. Embo Journal 17, 2663–2676. 10.1093/emboj/17.9.2663.

86. Schmitt, C., von Kobbe, C., Bachi, A., Panté, N., Rodrigues, J.P., Boscheron, C., Rigaut, G., Wilm, M., Séraphin, B., Carmo-Fonseca, M., and Izaurralde, E. (1999). Dbp5, a DEAD-box protein required for mRNA export, is recruited to the cytoplasmic fibrils of nuclear pore complex via a conserved interaction with CAN/Nup159p. Embo Journal 18, 4332–4347. 10.1093/emboj/18.15.4332.

87. Weirich, C.S., Erzberger, J.P., Berger, J.M., and Weis, K. (2004). The N-terminal domain of Nup159 forms a β-propeller that functions in mRNA export by tethering the helicase Dbp5 to the nuclear pore. Molecular Cell 16, 749–760. 10.1016/j.molcel.2004.10.032.

88. Napetschnig, J., Kassube, S.A., Debler, E.W., Wong, R.W., Blobel, G., and Hoelz, A. (2009). Structural and functional analysis of the interaction between the nucleoporin Nup214 and the DEAD-box helicase Ddx19. Proceedings of the National Academy of Sciences of the United States of America 106, 3089–3094. 10.1073/pnas.0813267106.

89. Seth, M., Shirayama, M., Gu, W.F., Ishidate, T., Conte, D., and Mello, C.C. (2013). The CSR-1 Argonaute Pathway Counteracts Epigenetic Silencing to Promote Germline Gene Expression. Developmental Cell 27, 656–663. 10.1016/j.devcel.2013.11.014.

90. Wedeles, C.J., Wu, M.Z., and Claycomb, J.M. (2013). Protection of germline gene expression by the *C. elegans* Argonaute CSR-1. Developmental Cell 27, 664–671. 10.1016/j.devcel.2013.11.016.

91. Branon, T.C., Bosch, J.A., Sanchez, A.D., Udeshi, N.D., Svinkina, T., Carr, S.A., Feldman, J.L., Perrimon, N., and Ting, A.Y. (2018). Efficient proximity labeling in living cells and organisms with TurboID. Nature Biotechnology 36, 880–887. 10.1038/nbt.4201.

92. Niepielko, M.G., Eagle, W.V.I., and Gavis, E.R. (2018). Stochastic Seeding Coupled with mRNA Self-Recruitment Generates Heterogeneous *Drosophila* Germ Granules. Current Biology 28, 1872–1881. 10.1016/j.cub.2018.04.037.

93. Kaur, T., Raju, M., Alshareedah, I., Davis, R.B., Potoyan, D.A., and Banerjee, P.R. (2021). Sequence-encoded and composition-dependent protein-RNA interactions control multiphasic condensate morphologies. Nature Communications 12. 10.1038/s41467-021-21089-4.

94. Uebel, C.J., Agbede, D., Wallis, D.C., and Phillips, C.M. (2020). Mutator Foci Are Regulated by Developmental Stage, RNA, and the Germline Cell Cycle in Caenorhabditis elegans. G3 (Bethesda) 10, 3719–3728. 10.1534/g3.120.401514.

95. Jeske, M., Müller, C.W., and Ephrussi, A. (2017). The LOTUS domain is a conserved DEAD-box RNA helicase regulator essential for the recruitment of Vasa to the germ plasm and nuage. Genes & Development 31, 939–952. 10.1101/gad.297051.117.

96. Kubíková, J., Reinig, R., Salgania, H.K., and Jeske, M. (2021). LOTUS-domain proteins - developmental effectors from a molecular perspective. Biological Chemistry 402, 7–23. 10.1515/hsz-2020-0270.

97. Zhang, Y., Wang, K., Yang, K.L., Shi, Y.Y., and Hong, J.J. (2021). Insight into the interaction between the RNA helicase CGH-1 and EDC-3 and its implications. Scientific Reports 11. 10.1038/s41598-021-99919-0.

## References

1 Kim, H. et al. A Co-CRISPR Strategy for Efficient Genome Editing in *Caenorhabditis elegans*. Genetics 197, 1069–U1037, 10.1534/genetics.114.166389 (2014).

2 Placentino, M. et al. Intrinsically disordered protein PID-2 modulates Z granules and is required for heritable piRNA-induced silencing in the *Caenorhabditis elegans* embryo. Embo Journal 40, 10.15252/embj.2020105280 (2021).

3 Wan, G. et al. ZSP-1 is a Z granule surface protein required for Z granule fluidity and germline immortality in *Caenorhabditis elegans*. Embo Journal 40, 10.15252/embj.2020105612 (2021).

4 Sheth, U., Pitt, J., Dennis, S. & Priess, J. R. Perinuclear P granules are the principal sites of mRNA export in adult *C. elegans* germ cells. Development 137, 1305–1314, 10.1242/dev.044255 (2010).

